# The Evolution of Sex Chromosome Dosage Compensation in Animals

**DOI:** 10.1101/2020.07.04.187476

**Authors:** Jiabi Chen, Menghan Wang, Xionglei He, Jian-Rong Yang, Xiaoshu Chen

## Abstract

The evolution of sex chromosomes in the XY or ZW systems shall lead to gene expression dosage problems, as in at least one of the sexes, the sex-linked gene dose has been reduced by half. It has been proposed, most notably by Susumu Ohno for mammals, that the transcriptional output of the whole sex chromosome should be doubled for a complete dosage compensation. However, due to the variability of the existing methods to determine the transcriptional differences between *S*ex chromosomes and *A*utosomes (S:A ratios) in different studies, whether clade-specific results are comparable and whether there is a more general model to explain dosage compensation states remain unanswered. In this study, we collected more than 500 public RNA-seq datasets from multiple tissues and species in major clades (including mammals, birds, fishes, insects, and worms) and proposed a unified computational framework for unbiased and comparable measurement of the S:A ratios of multiple species. We also tested the evolution of dosage compensation more directly by assessing changes in the expression levels of the current sex-linked genes relative to those of the ancestral sex-linked genes. We found that in mammals and birds, the S:A ratio is approximately 0.5, while in insects, fishes and flatworms, the S:A ratio is approximately 1. Further analysis showed that the fraction of dosage-sensitive housekeeping genes on the sex chromosome is significantly correlated with the S:A ratio. In addition, the degree of degradation of the Y chromosome may be responsible for the change in the S:A ratio in mammals without a dosage-compensation mechanism. Our observations offer unequivocal support for the sex chromosome insensitivity hypothesis in animals and suggest that the dosage sensitivity states of sex chromosomes is a major factor underlying different evolutionary strategies of dosage compensation.

## Introduction

The evolution of sex is generally credited with great evolutionary importance (Smith and Maynard-Smith 1978). On the molecular and genomic level, the origination of sex chromosomes was a hallmark of the evolution of sex in animals. Two of the most studied types of sex chromosome systems in the animal kingdom are the XY system and the ZW system. Despite many differences (Ezaz, et al. 2006), the XY and ZW systems both consist a pair of heteromorphic sex chromosomes, one of which is usually highly degenerate (W and Y) unless it is evolutionarily young, making the sex chromosomes very different from their ancestors.

One critical difference between sex chromosomes and autosomes is the dose of genes that reside on them. If Y/W-homologous is degenerated or inactivated, the dose of sex-linked genes should be halved relative to autosomal genes in the heterogamete sex (males in the XY system and females in the ZW system) (Naurin, et al. 2010), so it should be compensated. This further introduces the problem of dose inequality between heterogametic sex and homogametic sex and is particularly harmful to dose-sensitive genes (such as members of protein complexes) (Papp, et al. 2003; Pessia, et al. 2014; Pessia, et al. 2012). Studies concerning the dosage compensation of sex-linked genes should therefore shed light on general rules governing genetics and development, and they have aroused long-standing attention in the field of molecular evolution (Gartler 2014). In the current study, we will focus on gene dosage compensation between sex chromosomes and autosomes, rather than dosage inequality between sexes.

Due to their importance, multiple models/mechanisms have been proposed for the evolutionary resolution of dosage compensation (Charlesworth 1996, 1978; Straub and Becker 2007). There could be either complete or incomplete dosage compensation. For the complete dosage compensation, gene expression is doubled in a chromosome-wide manner. One of the most influential theories regarding the complete dosage compensation was Ohno’s hypothesis (Ohno 1967), which proposed that the transcriptional output of the entire X chromosome was doubled during the evolution of the XY system in mammals. In the genomic era, Ohno’s hypothesis was initially supported by the microarray-based observation that the X chromosome to autosome gene expression ratio (X:A) approximately equaled 1 (Nguyen and Disteche 2006). However, since about 10 years ago, Ohno’s hypothesis has been challenged by RNA-seq-based transcriptional profiles (Chen and Zhang 2016; Julien, et al. 2012; Pessia, et al. 2012; Xiong, et al. 2010), proteomic data (Chen and Zhang 2015), and direct comparison between the current X-linked genes and the ancestral X-linked genes (He, et al. 2011; Lin, et al. 2012; Marin, et al. 2017). These studies have reported that X:A approximately equaled 0.5, i.e., no double expression of the X chromosome was observed. The debate continued as several groups found support for Ohno’s hypothesis (Deng, et al. 2011; Kharchenko, et al. 2011; Lin, et al. 2011). These contrasting conclusions appeared influenced by whether the ancestral X chromosome (proto-X chromosome, hereinafter denoted as <u>X</u>) was involved (Julien, et al. 2012; Lin, et al. 2012) and by specific analytical approaches (Jue, et al. 2013) (see the first and second sections of the Results). In addition to the intense debate on mammals, complete dosage compensation for the XY system has been observed, sometimes in a tissue- or stage-specific manner, in Diptera (fly and mosquito) (Jiang, et al. 2015; Joshi and Meller 2017; Nozawa, et al. 2014; Vicoso and Bachtrog 2015). Thus, it is still an open question whether complete dosage compensation is applicable to a wide range of species. However, there could be incomplete or partial dosage compensation, in which the up-regulation of sex-linked genes appears on a gene-by-gene basis. The incomplete dosage compensation was first documented in chickens (Ellegren, et al. 2007; Itoh, et al. 2007) and subsequently found in most ZW species (Gu, et al. 2017; Naurin, et al. 2011; Wolf and Bryk 2011; Wright, et al. 2015) and some XY species (Wheeler, et al. 2016; White, et al. 2015).

In this study, we gathered over 500 public RNA-seq datasets from multiple tissues of various organisms in major clades of animals (including mammals, birds, fishes, insects and worms) and proposed a unified computational framework for unbiased and comparable measurement of the transcriptional ratios between the *S*ex chromosome and *A*utosomes (S:A ratios, including X:A ratios and Z:A ratios) among multiple species. Our aim was to provide unambiguous answers to two major questions: (i) whether complete dosage compensation is applicable to all mammals and to other animals, and (ii) whether there are factors other than sex chromosome degeneration underlying the evolution of complete *versus* incomplete dosage compensation.

## Results

### Construction of an extensive transcriptome dataset for assessment of sex chromosome dosage compensation

To enable an accurate assessment of dosage compensation among multiple species, we constructed an extensive transcriptome dataset from public RNA-seq datasets. To ensure sufficient data quality while maximizing the number of species and tissues, we selected the transcriptome dataset through two key steps: species filtering and expression screening. For species filtering, we chose species whose genomes were sequenced and annotated, with well-defined heteromorphic sex chromosomes (see Methods). For expression screening, we collected, from GEO/SRA, Illumina platform-based RNA-seq raw sequencing data with clear sex and tissue information, excluding pathological, stressed, genetically modified or unmappable (mapping rate for the short reads < 50%) samples. We only considered the top 20 most frequently studied tissues in each species and chose the one SRA record with the greatest sequencing depth for each tissue in each sex of each species (see Methods). The final transcriptome dataset consisted of 535 SRA files (Supplemental Table S1) from 32 species (Supplemental Table S2).

The transcript abundance in each sample was then estimated through a unified computational pipeline to ensure comparability (see Methods). Our estimated expression levels were highly correlated with previous RNA-seq and qRT-PCR reports (Xiong, et al. 2010), supporting the accuracy of our pipeline (Supplemental Fig S1).

### Choice of gene set for unbiased estimation of the S:A ratio

One major controversy surrounding evidence supporting/against Ohno’s hypothesis in mammals was how should filter the appropriate gene sets for fair expression comparisons between sex-linked genes and autosomal genes (Castagne, et al. 2011; Catalan, et al. 2018; Gu and Walters 2017; Jue, et al. 2013). One strategy, “filter-by-fraction” (Deng, et al. 2011), was to choose the identical fraction of highly expressed genes from the sex chromosomes and autosomes^9,13,14^. Another strategy, “filter-by-expression”, used a single expression cutoff value to select “expressed” genes on both the sex chromosomes and autosomes (Deng, et al. 2011; Kharchenko, et al. 2011; Lin, et al. 2011). To check whether these two strategies provide an unbiased estimation of the S:A ratio, we used an RNA-seq-based transcriptional profile of female human liver (Supplemental Table S3) to construct artificial test sets reflecting the models of complete or incomplete dosage compensation between the sex chromosome and the autosomes.

The actual autosomal gene expression levels in the liver were first used as the autosomal expression levels shared by the two models (Fig 1A-D, small red dots). A randomly chosen subset of autosomal genes, containing the same number of human X-linked genes, was designated as fake X-linked genes. The expression of these fake X-linked genes and the autosomal genes thus constituted a dataset that reflected the complete dosage compensation model (Fig 1A and C, small blue dots). Another dataset for the incomplete dosage compensation model was constructed by halving the expression level of the fake X-linked gene (Fig 1B and D, small blue dots). The S:A (or X:A in this specific case) ratios were then calculated for these two datasets by first applying either the filter-by-fraction (Fig 1A and B) or filter-by-expression (Fig 1C and D) strategy, followed by dividing the median expression level of the fake X-linked genes by that of the autosomal genes.

**Figure 1.**
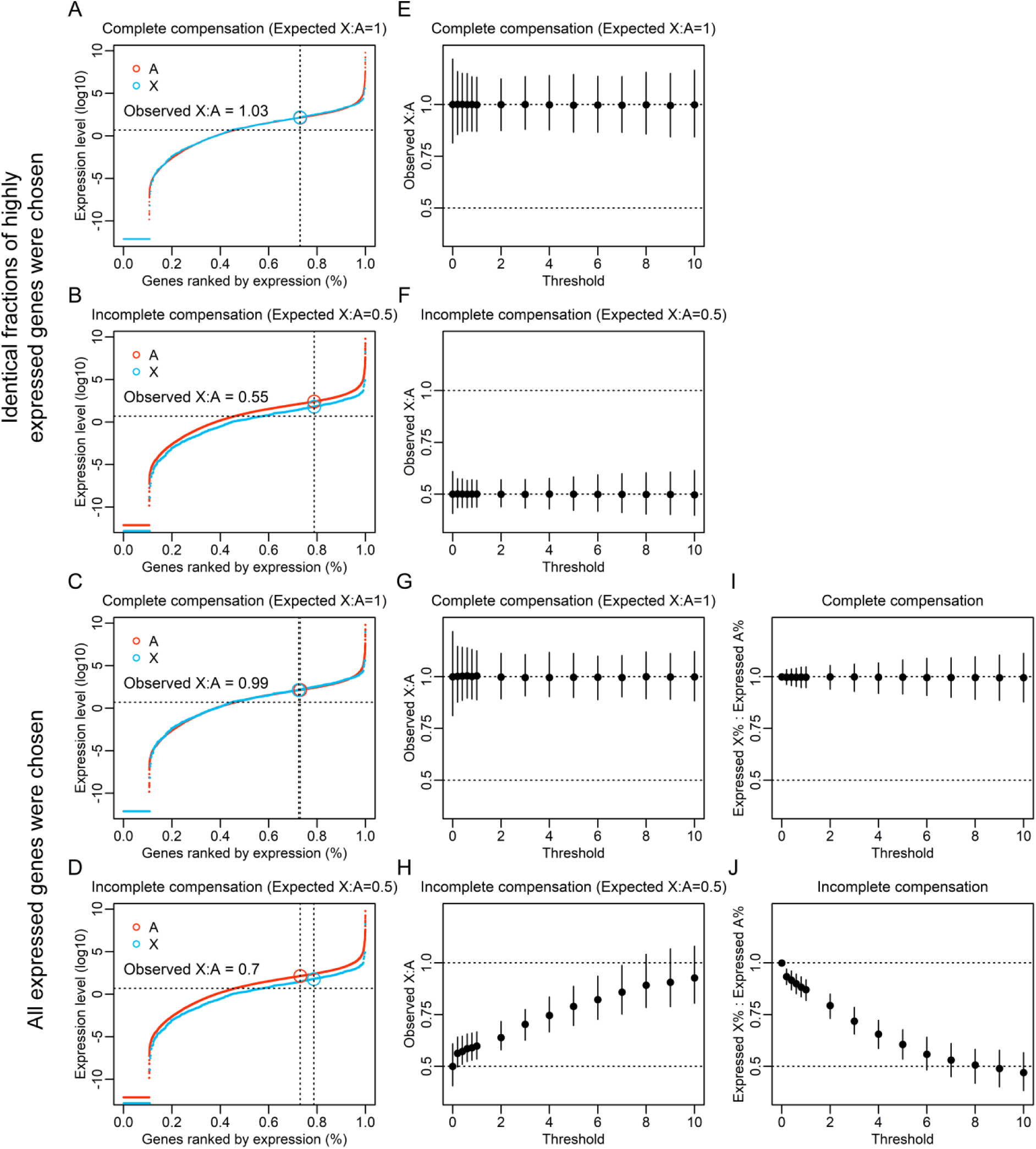
The impact of the gene-filtering strategy on assessing the complete dosage compensation between the sex chromosomes and autosomes. **(A)** The expression levels of autosomal genes in female human livers were converted with base-10 logarithmic transformation and ranked (red dots). In the complete dosage compensation model, autosomal genes with identical numbers of actual human X-linked genes were randomly selected as X-linked genes (blue dots). The median expression levels of the X-linked genes (blue circle) and the autosomal genes (red circle) were calculated by the filter-by-fraction strategy, in which the identical fraction of genes whose expression levels exceeded a certain threshold (FPKM > 2) was chosen. The threshold was indicated by the dotted horizontal line. **(B)** Similar to (A) except that in the incomplete dosage compensation model, the expression levels of the randomly picked X-linked genes were halved (blue dots). **(C)** Similar to (A) except that the median expression levels of the X-linked genes (blue circle) and the autosomal genes (red circle) were calculated by the filter-by-expression strategy, in which all genes with expression levels above a certain threshold were chosen. **(D)**. Similar to (C) except that in the incomplete dosage compensation model, the expression levels of the randomly picked X-linked genes were halved (blue dots). **(E-H)** Respectively corresponding to (A-D), the X:A ratio was calculated with different thresholds (FPKM > 0, 0.2, 0.4, 0.6, 0.8, 1, 2, 3, 4, 5, 6, 7, 8, 9, 10) and for different models by different strategies. The error bars represent the 95% confidence intervals for the range of X:A ratios assessed by 1,000 different random sets of X-linked genes picked from the autosomes. (**I** and **J**) Corresponding to **(G)** and **(H)**, the ratio of the fraction of expressed genes on the X chromosome and that on autosomes was calculated with different thresholds. The value of FPKM, representing the expression level, was converted with a base-10 logarithmic transformation. The expression levels of genes with FPKM values equal to 0 were set to one-tenth of the minimum non-zero expression abundance in each corresponding sample.

We first used a threshold of FPKM (*f*ragments *p*er *k*ilobase of exon per *m*illion fragments mapped) > 2 (which was previously used to find support for complete dosage compensation (Deng, et al. 2011), and see the next paragraph for other thresholds) to compare the performance of the two gene-filtering strategies under the complete/incomplete dosage compensation model. Specifically, according to the filter-by-expression strategy, genes with FPKM > 2 were directly included for calculating the X:A ratio. For the filter-by-fraction strategy, the fraction of genes with FPKM > 2 was calculated separately for the sex chromosome and autosomes, and the smaller fraction between these two was used as the fractional cutoff to choose the most highly expressed genes from both the X chromosome and autosomes. As a result, we found that the filter-by-fraction strategy provided unbiased estimations of the X:A ratio in both the complete dosage compensation and the incomplete dosage compensation models (Fig 1A and B). However, the filter-by-expression strategy was only accurate for the complete dosage compensation model (Fig 1C), while it overestimated X:A in incomplete dosage compensation model by ∼40%, from the expected X:A = 0.5 to the observed X:A ∼ 0.7 (Fig 1D).

To further examine the accuracies of the two gene-filtering strategies, we repeated the above analysis 1,000 times for each of the various thresholds up to an FPKM of 10 (only ∼20% of the genes met the criteria of FPKM > 10). We found that the filter-by-fraction strategy consistently enabled unbiased estimation of the X:A ratio regardless of the presence or absence of complete dosage compensation (Fig 1E and F). In contrast, the filter-by-expression strategy provided an accurate estimation of the X:A ratio in the complete dosage compensation model (Fig 1G), but not in the incomplete dosage compensation model (Fig 1H). As the FPKM threshold increased, the observed X:A ratio gradually increased and approached 1, significantly deviating from the expected value of 0.5 in the absence of complete dosage compensation (Fig 1H). These results showed that the filter-by-expression strategy is inherently biased toward complete dosage compensation, which is consistent with previous reports (Castagne, et al. 2011; Gu and Walters 2017; He, et al. 2011). It is worth noting that such an increase in the X:A ratio when a higher expression threshold was applied is very similar to the observation made in previous reports supporting Ohno’s hypothesis (Deng, et al. 2011; Sangrithi, et al. 2017), therefore casting doubts over their conclusions. In addition, we stress that observations of X:A in other studies larger than our simulated result (Fig 1H) were not necessarily support for globally increased expression on the sex chromosome, but could instead be explained by partial compensation derived from a subset of genes on the sex chromosome (Ellegren, et al. 2007; Lin, et al. 2012; Pessia, et al. 2012).

To better understand the source of bias of the filter-by-expression strategy, we calculated the fractions of genes expressed (FPKM exceeding a certain threshold) in the sex chromosome and the autosomes for various FPKM thresholds and plotted the ratio of these two fractions. The results indicated that without complete dosage compensation, an increase in the expression threshold resulted in over-exclusion of weakly expressed X-linked genes (Fig 1J) such that the median expression levels of X-linked genes were overestimated relative to those of autosomal genes. Note that such an excessive exclusion of X-linked genes occurred only in the absence of complete dosage compensation (Fig 1J); it did not occur when the dosage was completely compensated (Fig 1I), which again shows the inherent bias of the filter-by-expression strategy toward Ohno’s theory.

### The sex chromosome to autosome expression ratios in multiple species

Based on the filter-by-fraction strategy (same as below, unless otherwise stated), we compared the expression levels between sex-linked and autosomal genes in up to 20 tissues for each species. Our focus was on the sex chromosome dosage compensation relative to autosomes, rather than dosage inequality between sexes. We also did not attempt to identify the small subset of compensated genes that potentially exist. Specifically, regardless of the tissues or sex, if the species develops a fully functional mechanism for complete dosage compensation, we would observe S: A ∼ 1 (unless Y/W degeneration is at an early stage). In contrast, if the species has no dosage compensation at all, S: A ∼ 0.5. We stressed that incomplete dosage compensation should be assumed when S:A ∼ 1 can be statistically rejected, regardless the statistical significance of tests regarding S:A ∼ 0.5, which was just presented here as a reference model of RNA ratio equals DNA ratio. In addition, we collected the comparison results of the expression levels between sex-linked genes and autosomal genes of specific tissues/sex^38,45^ for future studies.

Our analysis revealed large variations in the levels of sex chromosome dosage compensation (FPKM > 0, maximizing the number of genes analyzed; Fig 2. For detailed results for each sex, see Supplemental Fig S2. For the statistical significance of each tissue, see Supplemental Table S4.) with considerable deviation from the phylogeny, suggesting multiple independent evolutionary origins of complete dosage compensation.

**Figure 2.**
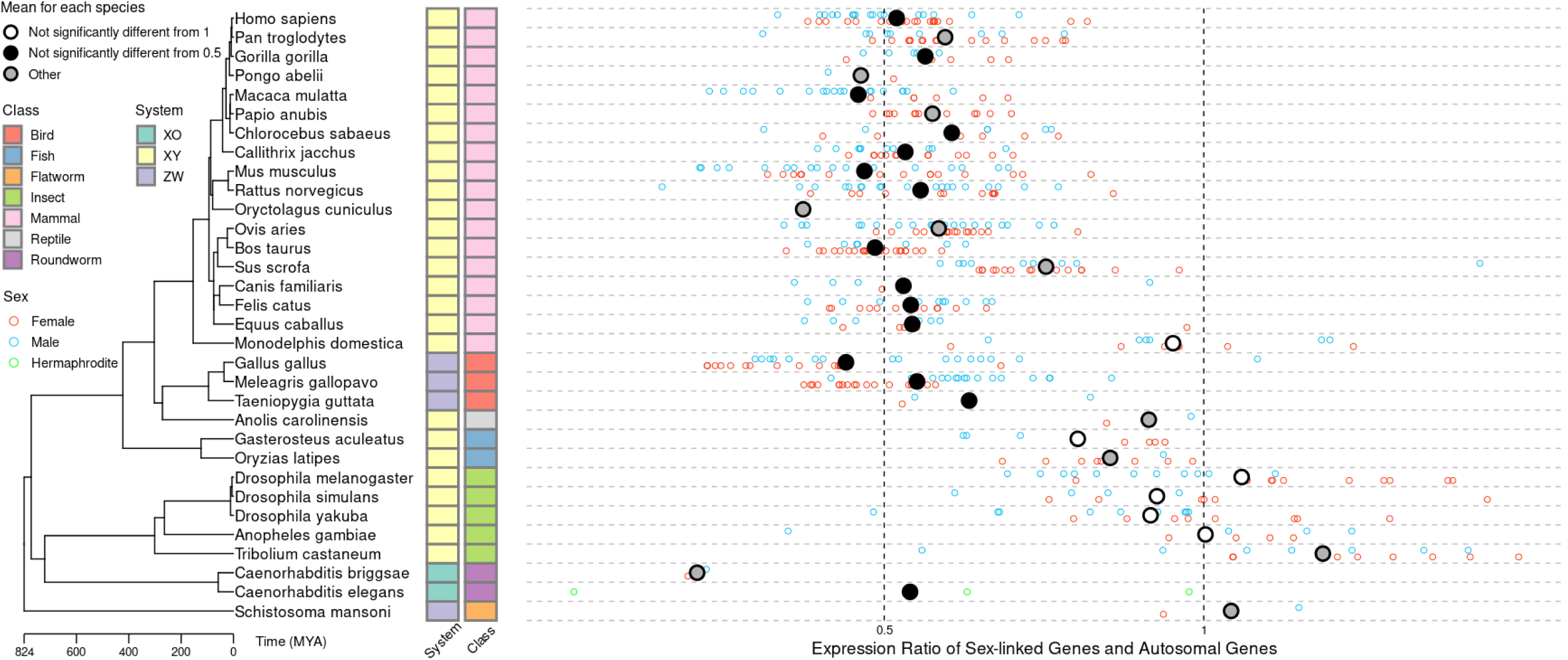
The sex chromosome to autosome expression ratios in multiple species revealed large variations in the level of sex chromosome dosage compensation that is apparently independent of their phylogeny. Shown on the left is a phylogenetic tree constructed for 32 species whose class and sex systems are labeled with colored blocks. The sex chromosome to autosome expression ratio (FPKM >0) for each tissue in each sex of each species is shown on the right. The red circles represent female tissues, the blue circles represent male tissues, and the green circles represent the hermaphrodite tissues. The average expression ratio of sex chromosome to autosomes for each species is indicated by a large circle, which was tested for equivalence to 1 or 0.5 by Student’s t test. Values not significantly different from 1 are marked with white circles, values not significantly different from 0.5 are marked with black circles, and other values are marked with gray circles.

Notably, in the clade of mammals, S:A was approximately 0.5 (Fig 2), confirming our previous results (Chen and Zhang 2016; Lin, et al. 2012; Xiong, et al. 2010) and those of another groups (Julien, et al. 2012; Pessia, et al. 2012). This conclusion is insensitive to the choice of expression threshold (Supplemental Fig S3). In particular, the S:A ratio of the opossum (*Monodelphis domestica*) was not significantly different from 1, suggesting that some subordinate taxa of mammals may have evolved complete dosage compensation, as supported by previous reports (Julien, et al. 2012; Lin, et al. 2012).

We also found a lack of complete dosage compensation in the bird clades, i.e., the S:A ratio was approximately equal to 0.5 (Fig 2). Although this observation was consistent with previous claims of incomplete dosage compensation (Mank 2013), it was quantitatively different from some other reports where the S:A ratio was found to be significantly greater than 0.5 (Brawand, et al. 2011). To clarify this inconsistency, we replicated the finding that the S:A ratio is significantly greater than 0.5 (Julien, et al. 2012; Lin, et al. 2012) using the expression levels of chicken genes provided in a previous report (Brawand, et al. 2011) (Supplemental Fig S4). However, a re-estimation of the expression levels through our pipeline using the raw sequencing data from the same report (Brawand, et al. 2011) yielded an S:A ratio of approximately 0.5 (Supplemental Fig S4), consistent with the conclusion obtained with our current dataset. As the current software and gene annotations used to calculate expression levels are more accurate than those used in previous reports, our current expression levels are most likely better than previously reported expression levels for accurate assessment of dosage compensation in birds (see Discussion). Nevertheless, our dataset and methods still yielded a conclusion consistent with previous findings that the S:A ratio is higher in males than in females (Supplemental Fig S2) (Julien, et al. 2012).

Some other specific cases are worth highlighting here. The green anole (*Anolis carolinensis*) is the only vertebrate that has been confirmed to support Ohno’s dosage compensation hypothesis (Marin, et al. 2017). In our dataset, we reached a similar conclusion that the S:A is close to 1 (∼ 0.9), although data from only two tissues were available for this study (Fig 2, see Methods).

With regard to fishes, the Japanese rice fish (*Oryzias latipes*) is a powerful model system for exploring the evolution of vertebrate sex chromosomes because it has XY sex chromosome pairs at relatively early stages of differentiation (Kondo, et al. 2006; White, et al. 2015). Before further degeneration of Y, the deviation of S:A from 1 is expected to be negligible, which was indeed observed (Fig 2). Similar observations have been found in the three-spined stickleback (*Gasterosteus aculeatus*), although its evolutionary history is somewhat different from the Japanese rice fish (Schultheiss, et al. 2015; White, et al. 2015) (Fig 2).

The gene dosage imbalance in flies created by hemizygosity of the X chromosome in males is compensated for by recruitment of the male-specific lethal (MSL) complex to X-linked genes and modification of chromatin (Jiang, et al. 2015; Joshi and Meller 2017; Mank 2013; Nozawa, et al. 2014). These findings were also confirmed by the observation from our unified computational pipeline, in which the S:A ratio was not significantly different from 1 (Fig 2).

In *Caenorhabditis elegans*, it was previously recognized that while a chromosome-wide balance of gene dosage is observed between sexes, no X-chromosome-wide mechanism exists to balance transcription between X and autosomes, as evidenced by the halved expression of X-linked relative to autosomal transgenes (Wheeler, et al. 2016). From our own analysis, we indeed observed the lack of complete dosage compensation for adult roundworms, despite the fact that each species contained only a few available transcription profiles (Fig 2, Supplemental Table S4). However, using only whole body tissue, we found no significant expression differences between the sex chromosome and autosomes in either female or male *Schistosoma mansoni* (Fig 2, Supplemental Table S4), indicating complete dosage compensation in flatworms. These findings are inconsistent with a report that female heterotypic parasites lack dosage compensation (Vicoso and Bachtrog 2011) but are consistent with some more recent reports (Picard, et al. 2018; Picard, et al. 2019).

To perform a more precise comparison excluding the brain and sex-related tissues (Wright, et al. 2012; Zimmer, et al. 2016), ten common tissues from the 32 species were selected to calculate the expression ratio of the sex chromosome to autosomes (Supplemental Fig S5, FPKM > 0). Consistent with the conclusions obtained from the expression ratios in up to 20 tissues per species, most mammals, birds and roundworms did not show complete dosage compensation. However, complete dosage compensation could be found in opossums, fishes, insects and flatworms, even though some species only contain two available transcription profiles. These results indicate that the complete dosage compensation of sex chromosomes has limited applicability in animal evolutionary history and has multiple independent origins.

We also replicated the above analysis using the filter-by-expression strategy. We noticed that when the depletion of expressed genes on sex chromosomes relative to autosomes became more dramatic when the expression threshold increased (Supplemental Fig S6, FPKM > 0; Supplemental Fig S7, FPKM > 2). This pattern exactly recapitulated the observation made with the filter-by-expression strategy from the aforementioned artificial dataset with fake X-linked genes, therefore suggesting that these transcriptional profiles conformed to the incomplete dosage compensation model (Fig 1J). In addition, larger S:A ratios were observed when the expression threshold increased (Supplemental Fig S8, FPKM > 0; Supplemental Fig S9, FPKM > 2), altering the conclusion of incomplete dosage compensation to complete dosage compensation, which was similar to previous reports in mammals (Deng, et al. 2011). This result again indicated the inherent bias of the filter-by-expression strategy toward complete dosage compensation.

### Sex to proto-sex expression ratios in different species

Ohno wrote in his seminal work that “During the course of evolution, an ancestor to placental mammals must have escaped a peril resulting from the hemizygous existence of all the X-linked genes in the male by doubling the rate of product output of each X-linked gene” (Ohno 1967). In other words, at least according to Ohno’s understanding, the complete dosage compensation should reflect the proposition of current sex-linked genes and ancestral autosomal genes that have evolved into sex chromosomes (proto-sex chromosomes, hereinafter denoted as <u>S</u>). Nevertheless, due to the inability to obtain gene expression levels on proto-sex chromosomes, previous attempts to verify Ohno’s hypothesis (S:<u>S</u> ∼ 1) were usually based on S:A (Deng, et al. 2011; Kharchenko, et al. 2011; Lin, et al. 2011), which implicitly assumed similar expression levels between <u>S</u> and ancestral autosomes (proto-autosomes, hereinafter denoted as <u>A</u>), as well as between A and <u>A</u>. These assumptions were explicitly examined in more recent studies, which directly evaluated S:<u>S</u> (Gu, et al. 2017; Nozawa, et al. 2014).

In sought of estimating S:<u>S</u> using our dataset, we used out-group species to identify proto-sex chromosomes and proto-autosomes via a previously described method (Julien, et al. 2012; Lin, et al. 2012). Humans and chickens have been identified as out-groups for many species (Lin, et al. 2012). In addition, *Drosophila melanogaster* and *Tribolium castaneum*, the transcriptional profiles of which are well enriched, are also suitable as out-groups for intra-tissue comparisons. After validating the suitability of these species as out-groups with a previous method (Pease and Hahn 2012) (see Methods, Supplemental Table S2), we were able to define <u>S</u> and <u>A</u> genes (see Methods, Supplemental Table S5). For example, a chicken autosomal gene with a one-to-one ortholog of a human X-linked (or A) gene is denoted as a human <u>S</u> (or <u>A</u>) gene.

We examined the similarity of paired tissues between each species and its out-group and found that the expression correlations of almost all paired tissues were greater than 0.5 (see Methods), supporting the reliability of this dataset. We thus used all paired tissues for subsequent analysis, which contained 230 pairs of tissue samples from 26 species (Supplemental Table S6). We examined the two assumptions regarding the expression levels of <u>S</u> and <u>A</u>. For some species, there were significant differences in the expression levels between the proto-sex-linked genes and proto-autosomal genes (Supplemental Fig S10). Additionally, the expression levels of proto-autosomal genes significantly differed from those of current autosomal genes in several species (Supplemental Fig S11). We then estimated the expression change from the proto-sex-linked genes to the current sex-linked genes by dividing S/<u>S</u> with A/<u>A</u>, thereby avoiding the assumptions of <u>S</u> = <u>A</u> and A = <u>A</u> (see Methods).

We discovered significant variations in the S to <u>S</u> expression ratios of the 26 species (Fig 3, Supplemental Table S6). In particular, mammals and most birds did not have complete dosage compensation, while fishes, insects and flatworms were completely dosage compensated. Overall, these observations are highly consistent with our results regarding the expression ratio between sex chromosomes and autosomes. Notably, results for some species are not available, such as roundworms and reptiles, because of the lack of tissues that were paired with the out-group (see Methods); results from some other species, such as the zebra finch (*Taeniopygia guttata*), were based on only one sex, which should be interpreted with caution. It should be emphasized that when we used the chickens as an out-group, we found that the S:<u>S</u> ratio of opossums was approximately 0.5, which changed the conclusion of complete dosage compensation derived from S:A to incomplete dosage compensation (see Discussion).

**Figure 3.**
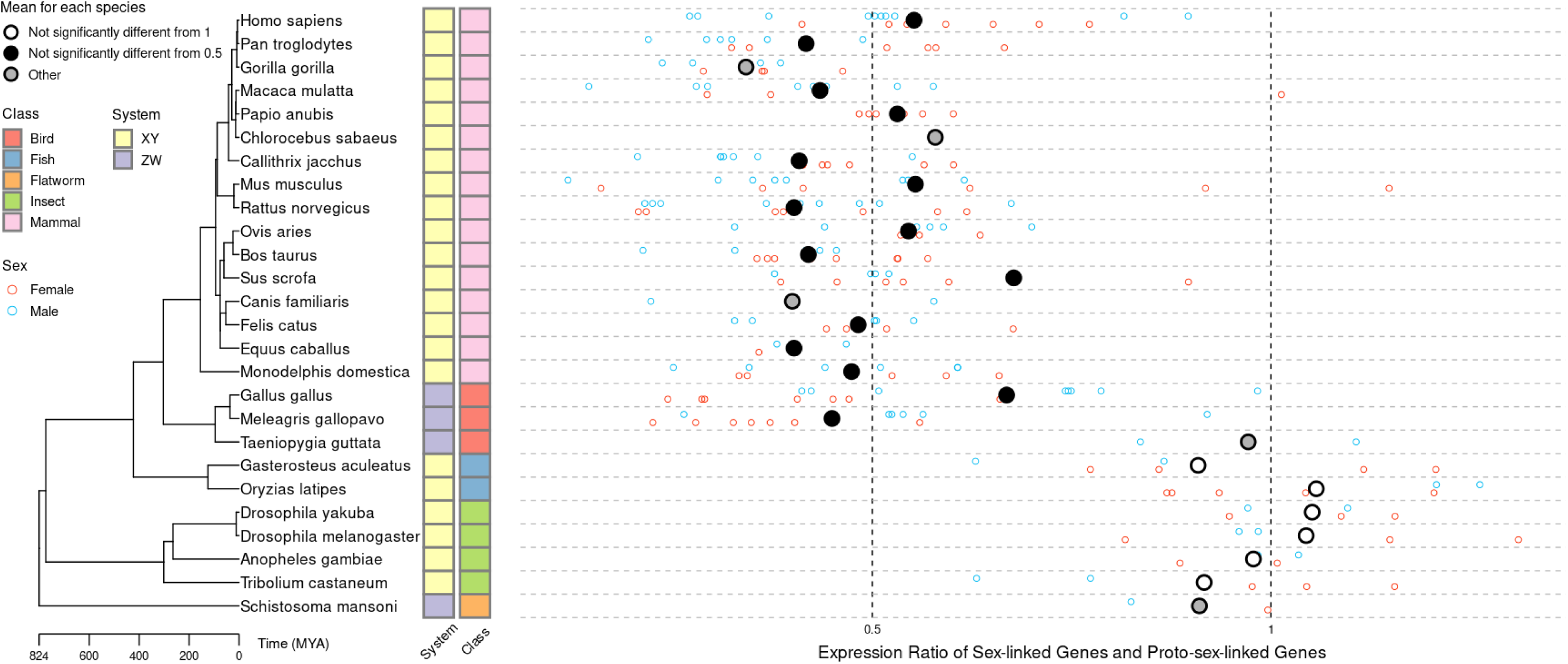
Significant variations in the expression ratios between the current sex-linked genes and the ancestral sex-linked genes. Shown on the left is a phylogenetic tree constructed for 26 species whose class and sex systems are labeled with colored blocks. The expression ratio between the current sex-linked genes and the ancestral sex-linked genes for each tissue in each sex of each species is shown on the right, with the red circles representing female tissues and the blue circles representing male tissues. The average expression ratio between the current sex-linked genes and the ancestral sex-linked genes in each species is indicated by a large circle, which was tested for equivalence to 1 or 0.5 by Student’s t test. Values not significantly different from 1 are marked with white circles, values not significantly different from 0.5 are marked with black circles, and other values are marked with gray circles.

Regardless, the above results systematically reflect the dosage differences between the sex chromosome and the autosomes among multiple species, again highlighting the variation in dosage compensation across different species.

### Selective pressure for complete dosage compensation varies with the fraction of dosage-sensitive genes

The above results and those of previous reports with unbiased estimation of S:A (Chen and Zhang 2016; Julien, et al. 2012; Lin, et al. 2012; Pessia, et al. 2012) have largely rejected the generality of complete dosage compensation for sex chromosomes, especially Ohno’s hypothesis for mammals. Instead, an alternative scenario, which we previously proposed as the “insensitive sex chromosome” hypothesis (Yang and Chen 2019), has risen. Specifically, we hypothesized that the sex (or the proto-sex) chromosomes tend to be depleted of dosage-sensitive genes; therefore, the selective pressure for complete dosage compensation was largely negligible (Pessia, et al. 2012; Yang and Chen 2019). Notably, our hypothesis was congruent with previous findings that the few genes retained by the mammalian Y and avian W chromosomes tend to be dosage-sensitive (Bellott, et al. 2014; Bellott, et al. 2017).

To assess the validity of the insensitive sex chromosome hypothesis, we examined the potential depletion of dosage-sensitive genes on sex chromosomes. To this end, we identified housekeeping genes, which are widely considered as dosage-sensitive (Bar-Even, et al. 2006; Newman, et al. 2006), and checked their distribution on autosomes and sex chromosomes (see Methods, Supplemental Table S7). Consistent with the insensitive sex chromosome hypothesis, we found a lack of housekeeping genes on the sex chromosomes of mammals and birds (Fig 4, middle). However, for fishes and insects, the fractions of housekeeping genes found on sex chromosomes and autosomes were similar (Fig 4, middle). More importantly, we noticed that the S:A ratio increased as the fraction of housekeeping genes on the sex chromosomes relative to that on the autosomes increased (Fig 5A), indicating that the evolutionary constraint on complete dosage compensation is stronger when the fraction of sex-linked dosage-sensitive genes is higher. This result remained unchanged when the phylogenetic interdependence was controlled by phylogenetic independent contrast analysis (Stone, et al. 2011) (Supplemental Fig S12). Note that, for some species, such as flatworms, we were unable to obtain a list of housekeeping genes due to the scarcity of tissues with transcriptional profiles.

**Figure 4.**
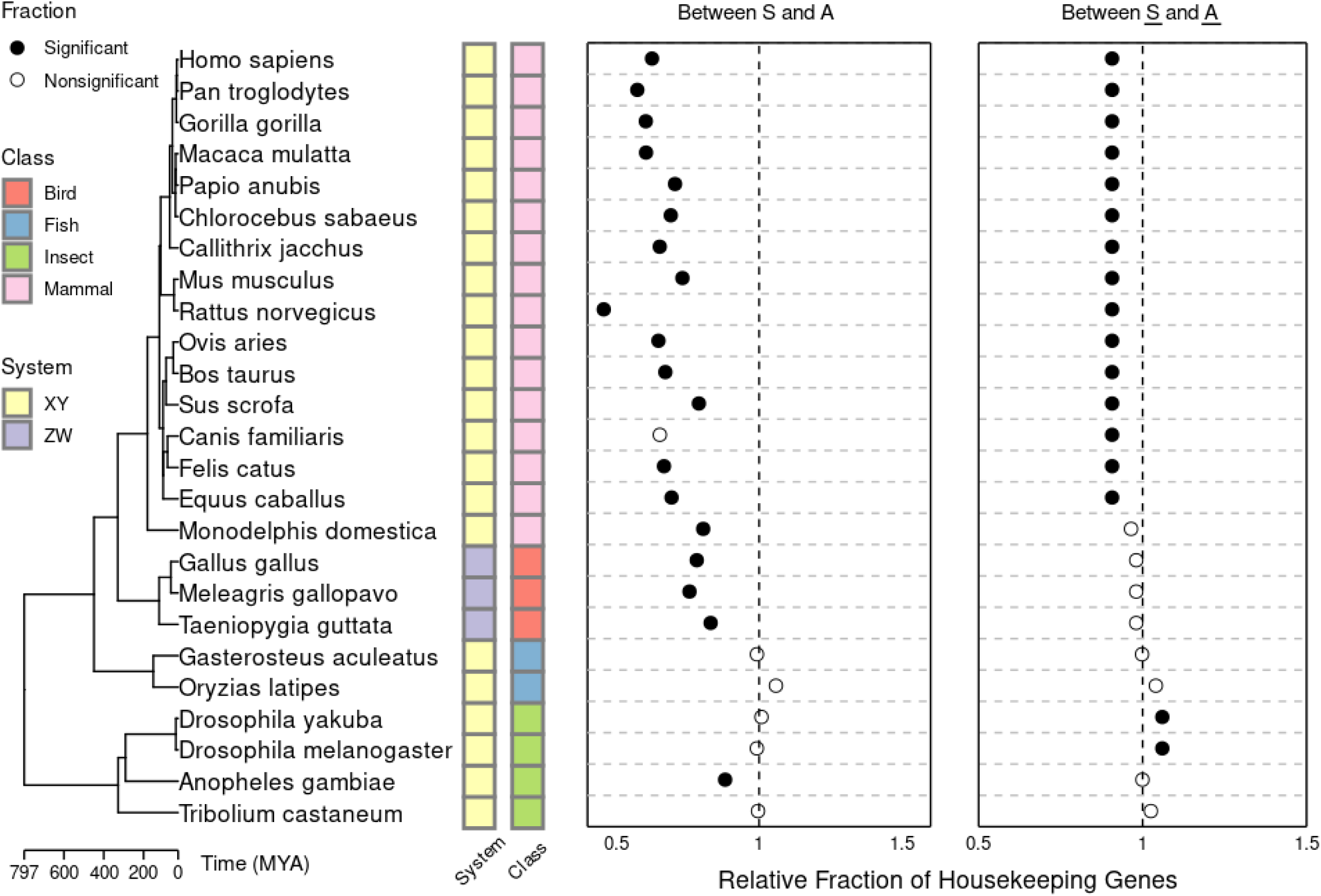
Significant variations in the relative fraction of housekeeping genes between sex chromosomes and autosomes in multiple species. Shown on the left is a phylogenetic tree constructed for 25 species whose class and sex systems are labeled with colored blocks. The fraction of housekeeping genes on the current sex chromosome relative to that on the current autosomes in each species is shown in the middle, and the fraction of housekeeping genes on the ancestral sex chromosome relative to that on the ancestral autosomes is shown on the right. The significance of differences in the relative fractions of the housekeeping genes was tested by the chi-square test. The black circles indicate a significant difference from 1, and the white circles indicate values similar to 1.

**Figure 5.**
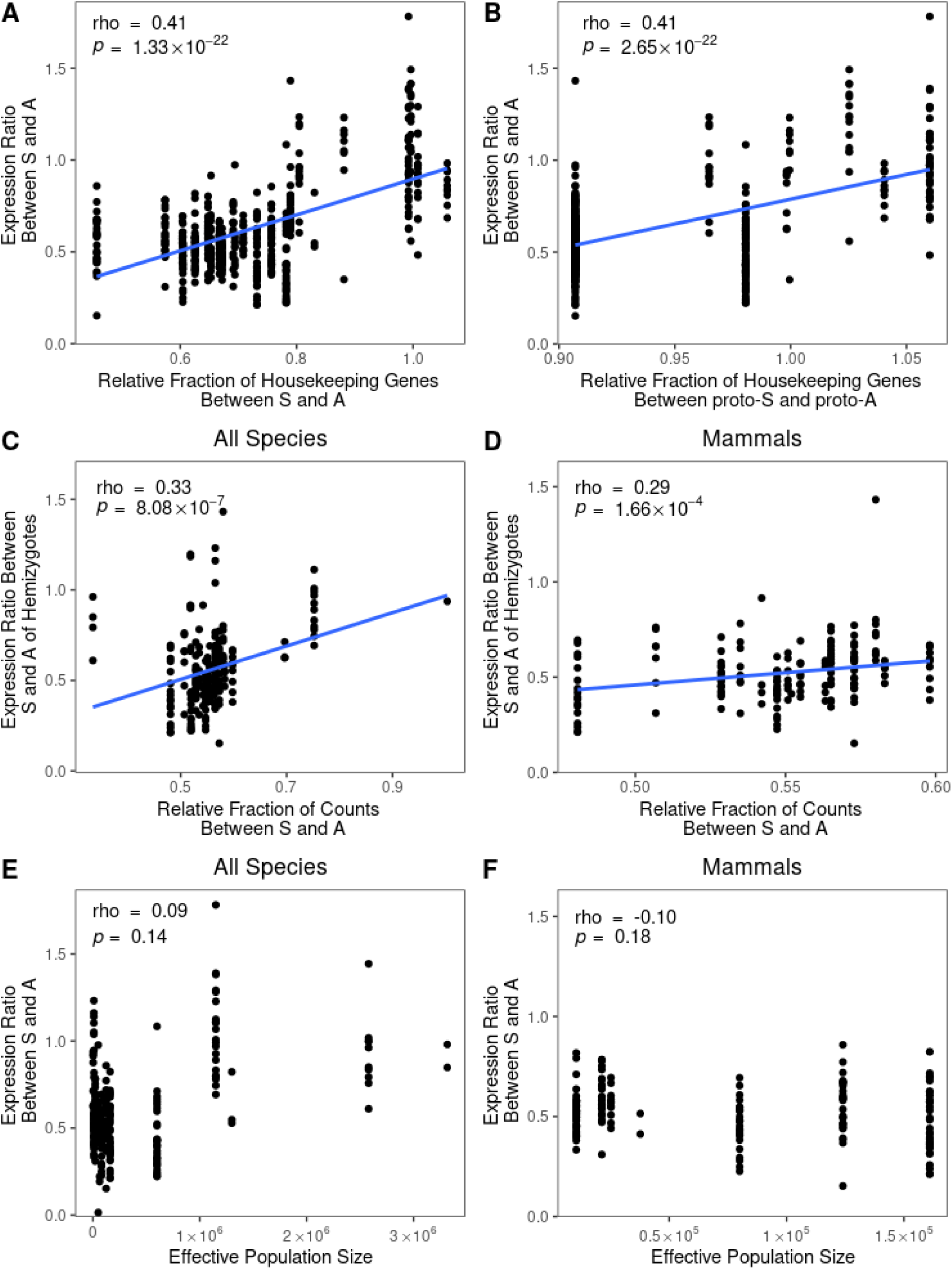
Possible factors underlying the large variations of the sex chromosome to autosome expression ratio. The fractions of housekeeping genes on the current sex chromosome relative to that on the current autosomes in each species **(A)** and between the ancestral sex chromosome and the ancestral autosomes **(B)** were significantly correlated with the sex chromosome to autosome expression ratios. The relative fraction of counts at the DNA level between sex chromosomes and autosomes was significantly correlated with the sex chromosome to autosomes expression ratios in all species **(C)** and in mammals **(D)**. The sex chromosome to autosome expression ratios were independent of the effective population size in all species examined **(E)** and in mammals **(F)**. Each dot represents the corresponding value in each tissue in each sex of each species.

The depletion of housekeeping genes on sex chromosomes relative to autosomes may have evolved through two nonexclusive mechanisms. The proto-sex chromosome may already be depleted with housekeeping genes (“insensitive proto-sex chromosome”). Alternatively, the housekeeping genes may have been gradually removed from the sex chromosome (“removal of dosage sensitive genes”). The removal of dosage-sensitive genes was previously supported by a few studies (Hurst, et al. 2015; Potrzebowski, et al. 2008), although the timing of gene migration from the X chromosome is much later than the time when common mammalian ancestors appeared (Potrzebowski, et al. 2008). We hereby tested the scenario of insensitive proto-sex chromosome (see Methods). As a result, we found that for all but one of the mammals, the proto-sex chromosomes contained a significantly smaller fraction of housekeeping genes relative to the proto-autosomes (Fig 4, right). Theoretically, species containing proto-sex chromosomes with fewer housekeeping genes should be under weaker natural selection for complete dosage compensation between sex chromosomes and autosomes. Our observations are therefore consistent with the general absence of complete dosage compensation for sex chromosome in mammals. Conversely, the proto-sex chromosomes of insects have more housekeeping genes than those of the proto-autosomes (Fig 4, right), which likely resulted in greater natural selection for (and therefore the evolution of) complete dosage compensation. Note that a few species with S:A ratios of approximately 0.5, such as birds, had similar fractions of housekeeping genes between proto-sex chromosomes and proto-autosomes, suggesting that the evolutionary dynamics of dosage compensation for such species have not been influenced by the distribution of dosage-sensitive genes in proto-chromosomes (Fig 4, right).

In addition, we examined the relationship between the S:A ratio and the fraction of housekeeping genes on proto-sex chromosomes relative to the proto-autosomes and found a significant positive correlation (Fig 5B). This result indicates that the evolutionary constraint on S:A complete dosage compensation is stronger if the fraction of housekeeping genes on proto-sex chromosomes is higher. Taken together, our findings suggest that in species without detectable sex chromosome complete dosage compensation, the sex chromosome evolved from a pair of (part of) autosomes with significant depletion of dosage-sensitive genes, thereby supporting the insensitive sex chromosome hypothesis.

### Y/W degeneration determines the sex chromosome to autosome expression ratio in heterogametic sex

Mammals are of common origin and contain the same nonsensitive genes on the proto-sex chromosome (Fig 4, right), but they have apparent variation in S:A (Fig 2 and 3). In the absence of complete dosage compensation, such variation in S:A should be explained by either partial compensation of individual genes or different levels of Y/W chromosome degeneration. More specifically, the S:A ratio will gradually decrease as the Y/W chromosome degenerates.

However, because Y/W chromosomes contain numerous repeat and amplicon sequences, most genome projects have excluded Y/W chromosomes from their analyses (Page, et al. 2010), making the level of Y/W chromosome degradation in each species difficult to quantify. To resolve this issue, we approximated the degree of sex chromosome degradation in 22 species by calculating the S:A ratio at the DNA level using DNA-seq data (see Methods). Due to the shortening of Y/W chromosomes, accumulation of mutations in Y/W-linked genes and other similar processes, some X/Z-linked genes lack homologues on the Y/W chromosome such that the S:A ratio at the DNA level will be less than 1 in males if the Y/W chromosome is (partially) degenerated. Consistent with this expectation, the S:A ratio at the DNA level (i.e., relative fractions of counts between S and A) was approximately 1 in human females and approximately 0.5 in human males (Supplemental Table S8). Subsequently, we estimated the degeneration of the Y/W chromosome by dividing the DNA-level S:A ratios of heterogametic sex divided by that of homogametic sex (see Methods).

Consistent with our hypothesis, the S:A expression ratios in the heterogametic sex were significantly positively correlated with the relative fractions of counts between S and A at the DNA level in all species (Fig 5C) and in mammals alone (Fig 5D). These results indicate that Y/W degeneration is an important factor affecting sex chromosome to autosome expression ratios, which is more important evidence supporting the hypothesis of insensitive sex chromosomes.

### Effective population size cannot explain the sex chromosome to autosomes expression ratio in all species

Effective population size (*N*_e_) of the examined species is another potential general factor that may explain the observed variation in S:A, as natural selection should be greater in species with large *N*_e_. We examined the relationship between *N*_e_ (Supplemental Table S9) and the S:A expression ratio among animals in several diverse phylogenetic groups covered by our dataset. As a result, we found no significant positive correlation between *N*_e_ and the S:A expression ratio in the species we tested (Fig 5E) or in mammals (Fig 5F). The above results indicate that the selection strength approximated by *N*_e_ cannot fully explain the variation of S:A among species, and other more important factors may underline the development of dosage-compensation mechanisms, such as distribution of dosage-sensitive genes between sex chromosomes and autosomes. Note, however, that our analysis did not consider the sex chromosome-specific *N*_e_, which was previously proposed as a reason underlying the more frequently observed complete dosage compensation for the XY system relative to the ZW system (Mank 2013; Mullon, et al. 2015). Additionally, the statistical power of this analysis may have been undermined by the limited number of species with estimated *N*_e_.

## Discussion

To assess the applicability of complete dosage compensation for sex chromosomes in a broad evolutionary context, we compiled a large transcriptome dataset for up to 20 tissues in 32 species representing the major clades of animals, including mammals, birds, reptiles, fishes, insects and worms. Using an unbiased computational method, we revealed various patterns of complete/incomplete dosage compensation across different species. We also tested the complete dosage compensation by using out-group species to compare changes in the expression levels of the current sex-linked genes relative to that of the proto-sex-linked genes. We found an S:A ratio of approximately 0.5 in mammals and birds and an S:A ratio of approximately 1 in insects, fishes and flatworms. Further analysis showed that the fraction of dosage-sensitive housekeeping genes on the sex chromosome was significantly correlated with the S:A ratio, and the fraction of housekeeping genes on the proto-sex chromosomes had a major effect on whether a species evolved a dosage-compensation mechanism. Furthermore, we found that the degree of Y or Z chromosome degeneration strongly influenced the S:A expression ratio. Our observations emphasized that the hypothesis of sex chromosome insensitivity can better describe dosage states of sex chromosomes in mammals and suggested that elucidation of the dosage-sensitivity properties of sex chromosomes can help reveal the different evolutionary strategies of dosage balancing between sex chromosomes and autosomes.

By a unified analysis of multiple species, we have described the evolution of complete dosage compensation, the results of which (Fig 2 and 3) are largely consistent with those of previous studies. Through simple comparison, we found that the expression levels of the entire sex chromosomes and autosomes of the Japanese rice fish are equivalent, verifying that its Y chromosome is newly formed (Kondo, et al. 2006). We also observed similar expression of sex chromosomes and autosomes in *Drosophila melanogaster* and *Drosophila yakuba*, corroborating previous reports of complete dosage compensation in Drosophila (Conrad and Akhtar 2012). In both sexes of mosquitos (*Anopheles gambiae*), the median expression ratios of X-linked genes to autosomal genes were close to 1, which was the same as previous results (Rose, et al. 2016). In beetles (*Tribolium castaneum*), we observed greater expression of X-linked genes than that of autosomal genes in both males and females, consistent with the finding of a previous study (Prince, et al. 2010) (but also see (Gu and Walters 2017; Mahajan and Bachtrog 2015)). We also found that Z-linked expression in *Schistosoma mansoni* was comparable to autosomal expression in females and males, which is consistent with complete dosage compensation (Picard, et al. 2018; Picard, et al. 2019).

However, our results in mammals, along with those from other groups (Julien, et al. 2012; Lin, et al. 2012; Marin, et al. 2017; Pessia, et al. 2012; Xiong, et al. 2010), were inconsistent with Ohno’s hypothesis. Previous studies supporting Ohno’s theory in mammals (Deng, et al. 2011; Kharchenko, et al. 2011; Lin, et al. 2011) or complete dosage compensation in other species (Picard, et al. 2018) have used the filter-by-expression strategy, which biases toward the dosage-compensation model even if the complete dosage compensation was absent (Fig 1). Indeed, genes with clear expression-doubling mechanisms have never been found in mammals. We have thus proposed an alternative hypothesis for Ohno’s theory, namely, the insensitive sex chromosome hypothesis. We also verified the predictions of our hypothesis that the distribution of dosage-sensitive genes and Y/W chromosome degeneration both affect the expression differences between the sex chromosome and autosomes, thus providing an evolutionary explanation for the formation of different dosage states between the sex chromosomes and autosomes in different species.

Surprisingly, there was no dosage compensation in female chickens (Fig 2 and 3), even though we observed that the S:A ratio was higher in males than in females (Supplemental Fig S2). Previous studies have reported partial dosage compensation (Julien, et al. 2012; Lin, et al. 2012). Both Julien et al. and Lin et al. used the transcriptome dataset obtained by Kaessmann’s group in 2011 (Brawand, et al. 2011), and we successfully replicated their results with these datasets (Supplemental Fig S4). We believe that our conclusions are more accurate than theirs for several reasons. First, our current dataset consists of transcriptional profiles from different research groups that were rigorously screened, thereby decreasing the likelihood of similar errors. Second, the transcriptional profiles we selected have a considerably larger sequencing depth and, thus, can more accurately reflect the expression levels of highly expressed genes and weakly expressed genes compared to transcriptional profiles with less depth. Third, the genome sequences and genome annotations we used are relatively new and, thus, should be more accurate than the old versions of genome sequences and annotations. Fourth, the turkey (*Meleagris gallopavo*), which is closely related to chickens, was not observed to have complete dosage compensation either. However, further evidence, such as quantitative proteome data in multiple species, is needed to confirm the presence of incomplete dosage compensation in birds.

The dosage state of opossums is controversial. Julien et al. reported that, compared with orthologs in chickens, the X-linked genes of opossums had substantial or complete dosage compensation (Julien, et al. 2012), while Lin et al. showed very little X compensation (Lin, et al. 2012). We used our chicken transcriptome dataset to estimate the expression levels of ancestral genes and of all protein-coding genes with one-to-one orthologs in the chickens, and we observed an S:<u>S</u> ratio of approximately 0.5 (Fig 3), indicating no dosage compensation for the sex chromosome of opossums. In addition, we observed an S:A ratio of ∼1 (Fig 2), an <u>S</u>:<u>A</u> ratio of ∼1 (Supplemental Fig S10) and an A:<u>A</u> ratio <1 (Supplemental Fig S11). These results indicate that the autosomes of opossums, rather than the sex chromosome, are subjected to regulation for complete dosage compensation. In other words, the overall downregulation of autosomal expression compensates for the difference in expression between autosomes and X chromosomes due to Y chromosome degeneration.

The dosage state between sex chromosomes and autosomes across the 32 species in the dataset compiled here cannot be fully explained by Ohno’s hypothesis. We therefore proposed the novel hypothesis of sex chromosome insensitivity to resolve this apparent lack of a theoretical foundation for the evolution of dosage of sex chromosomes. Assuming dosage sensitivity of housekeeping genes, we showed that species with dosage-compensated sex chromosomes tended to have a higher fraction of dosage-sensitive genes on their sex chromosomes than other species, thereby lending support for sex chromosome insensitivity. It is worth noting that there were still a few exceptions, such as the dog (*Canis familiaris*), zebra finch and mosquito, which may be caused by the scarcity of tissues with transcriptional profiles in these species and the consequential difficulties in identifying housekeeping genes. This problem is expected to be overcome by the availability of more transcriptional profiles. In addition, because housekeeping genes are also highly expressed genes, it was not possible in this study to discriminate whether uncompensated ancestral genes were indeed insensitive or were overall transcribed at low levels. Distinguishing these two possibilities will require single-cell transcriptional profiles, with which expression noise can be measured for direct identification of dosage-sensitive genes.

## Methods

### Genomic, comparative genomic and transcriptomic data

The genomes of different animals were selected according to several criteria. Briefly, as of June 2017, each species had to have (i) chromosome-level genome assembly data with well-defined sex chromosomes and homolog (one-to-one ortholog) information available in EnsEMBL (www.ensembl.org), and (ii) RNA-seq transcriptome data for normal tissues with clear sex and tissue annotations in the GEO/SRA database. More details are described below.

Genomes with an assembly level of “chromosomes” and a contig ratio of 50% or less were selected from Ensembl. For each selected genome, the chromosome annotated as X/Z/Y/W was considered to be the sex chromosome. For genomes without this type of sex chromosome annotation, we assessed the reference articles to manually determine the sex chromosomes, such as LGb for the green anole lizard (*Anolis carolinensis*) (Rupp, et al. 2017), Group XIX for the three-spined stickleback (*Gasterosteus aculeatus*) (Bakker, et al. 2017), and chromosome 1 of the Japanese rice fish (*Oryzias latipes*) (Kondo, et al. 2006). Genomes were not considered for our study if we could not determine the sex chromosomes by the above procedure. Ultimately, 32 genomes were included as candidates for RNA-seq transcriptional profile selection (Supplemental Table S2). The genomic assembly, protein-coding gene annotation and homology (one-to-one orthologs) information for each selected species was downloaded from Ensembl version 89 and Ensembl metazoan version 36. The phylogenetic tree of these species was downloaded from NCBI Taxonomy.

To collect relevant RNA-seq transcriptional profiles, we downloaded a compressed SQL file named “SRAmetadb.sqlite” on June 2017, via the R package SRAdb (Zhu, et al. 2013). In addition, we used the Perl package “Bio::LITE::Taxonomy::NCBI” to find the taxonomic IDs of all subspecies of each selected species, thereby allowing the inclusion of the subspecies RNA-seq transcriptional profiles. Then, based on the metadata from the GEO or SRA entries, the transcriptional profiles of the species that have been chosen were further screened with the following criteria: (i) the transcriptional profiles were RNA-seq profiles based on the Illumina sequencing platform; (ii) the transcriptional profiles had clear sex and tissue information; (iii) the transcriptional profiles were not derived from pathological, stress-treated or genetically modified samples, and the transcriptional profiles of roundworms and chickens should be derived from adults (Deng, et al. 2011; Xiong, et al. 2010); (iv) the transcriptional profiles were among the top 20 most frequently studied tissues (the ten most commonly studied transcriptional profiles in cross-species tissues were also selected to enable tissue-matched comparative analysis among species); and (v) the SRA records had the highest sequencing depth for each tissue in each sex of each species. If the SRA record could not be downloaded or if more than 50% of the short reads could not be mapped to the corresponding genome, we discarded the SRA record and selected the next best record according to the above rules until all conditions were met. The final transcriptome dataset consisted of 535 SRA files (Supplemental Table S1) from 32 species.

We downloaded the SRA files using SRA Toolkit (Sherry 2000), generated short-read alignments with the corresponding genomes by HISAT2, and quantified expression with StringTie (Pertea, et al. 2015). The resulting FPKM value was used as the expression level for each gene (Supplemental Table S3). FPKM values were also transformed to *T*ranscripts *p*er *m*illion reads (TPM) values, a metric known to be better scaled for comparison between samples (Wagner, et al. 2012), to allow identification of housekeeping genes.

### Validation of the selected out-group species

A previous study on sex chromosome evolution found that humans (*Homo sapiens*) and chickens (*Gallus gallus*) can be used as out-groups for many species (Lin, et al. 2012). In addition, the fruit fly (*Drosophila melanogaster*) and the beetle (*Tribolium castaneum*) are also suitable candidate out-groups because transcriptional profiles are available for multiple tissues. We validated the suitability of these four species as out-groups for the other species with a previously described method that assesses the independent origins of sex chromosomes (Pease and Hahn 2012).

Briefly, for any pair of species, one was regarded as the focal species and the other was regarded as the candidate out-group. For all orthologous pairs of genes between the two species, the orthologous relationships were randomly shuffled regardless of their chromosomal location. This random process was repeated 1,000 times, resulting in an expected distribution of the number of genes for each chromosome in the candidate out-group that was orthologous to sex-linked genes in the focal species. This expected distribution was then compared with the actual number derived from the real orthologous relationship to obtain the degree of enrichment of each chromosome in the candidate out-group that was orthologous to sex-linked genes in the focal species. If a chromosome of the candidate out-group was significantly enriched (P ≤ 10^-3^) with genes orthologous to the sex-linked genes in the focal species, we considered this chromosome of the candidate out-group to be the proto-sex chromosome of the focal species. If the sex chromosome of the candidate out-group was not the proto-sex chromosome, the candidate was considered to be an appropriate out-group for the focal species because by definition, the proto-sex chromosome must be an autosome.

With regard to vertebrates, after confirmation using the above procedure, chickens were used as the out-group for mammals, and humans were used as the out-group for birds, reptiles and fishes. With regard to invertebrates, beetles formed the out-group for flies and mosquitos, and fruit flies formed the out-group for beetles and worms (Supplemental Table S2).

### Selection of transcriptional profiles for comparison between current genes and ancestral genes

We used expression data from the same tissue in the focal species and the out-group species to compare expression between the current and ancestral chromosomes. To further ensure the comparability of the expression levels of the paired tissues, we performed a Spearman correlation analysis on the logarithmic values of FPKM in the transcriptional profiles between the focal species and the out-group species. To avoid division by zero, genes with FPKM values of 0 were assigned to one-tenth of the minimum non-zero FPKM in the corresponding transcriptional profile. Transcriptional profiles with no paired data were discarded. Ultimately, 230 transcriptional profiles from 26 species were retained in the comparison of the expression between current and ancestral chromosomes (Supplemental Table S6).

### Expression ratio between sex-linked genes and autosomal genes

For an unbiased estimation of the S:A expression ratio, we adopted the filter-by-fraction strategy to choose an identical fraction of highly expressed genes from the sex chromosome and the autosomes in our analysis (see the second section of the Results). Specifically, if the fraction of genes on the X/Z chromosome that were greater than a certain expression threshold (defined as expressed genes) was less than that on autosomes, all these expressed genes on the X/Z chromosome were chosen; the most highly expressed autosomal genes were chosen based on the identical fraction in the X/Z chromosome, and vice versa. We then calculated the S:A expression ratio as the median expression level of the chosen sex chromosome genes divided by the median expression level of the chosen autosomal genes (see Formula 1 below). The 95% confidence intervals of the S:A ratios were estimated by bootstrapping the genes 1,000 times (Supplemental Table S4). In addition, we also applied the filter-by-expression strategy as a comparison with the filter-by-fraction strategy, i.e., all sex chromosomes and autosomal genes that passed a certain expression threshold were chosen. We then also used the same formula and statistical method of the filter-by-fraction strategy to calculate the S:A expression ratio and 95% confidence interval.

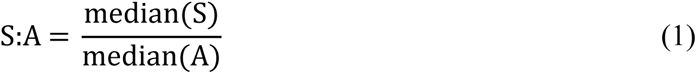

### Expression ratio between current sex-linked genes and ancestral sex-linked genes

Testing Ohno’s hypothesis by the estimation of S:A was based on two implicit assumptions. While the expression of ancestral sex chromosomes was the same as that of ancestral autosomes, the expression of ancestral autosomes was the same as that of current autosomes. However, it is possible that one or both of these assumptions were violated. As suggested by previous studies (Julien, et al. 2012; Lin, et al. 2012), direct testing of Ohno’s hypothesis on sex chromosome dosage compensation should involve comparison of the expression levels of current sex-linked genes (S) with those of the ancestral sex-linked genes (<u>S</u>) that existed before sex chromosome divergence.

We first defined the <u>S</u> and ancestral autosomal genes (<u>A</u>) in the predetermined out-groups via one-to-one orthologs of genes in the focal species. Similar to previous studies (Julien, et al. 2012; Lin, et al. 2012), a chicken autosomal gene was referred to as a human <u>S</u> (or <u>A</u>) gene if it was a one-to-one ortholog of a human X-linked (or A) gene (Supplemental Table S5). We emphasized that <u>S</u> genes were usually distributed in more than one autosome of the out-group species (Stiglec, et al. 2007).

We then evaluated the S and <u>S</u> expression ratios by a previously proposed method (Julien, et al. 2012; Lin, et al. 2012). Specifically, we first calculated the ratio of expression levels between each sex-linked gene and its one-to-one orthologous autosomal gene in the out-group. To avoid division by zero, genes with FPKM values of 0 were assigned to one-tenth of the minimum non-zero FPKM in the corresponding transcriptional profile. We then normalized the median S to <u>S</u> expression ratio by the ratio of the median expression of autosomal genes in the focal species to the median expression of autosomal genes in the out-group (see Formula 2 below). The 95% confidence intervals of these ratios were estimated by bootstrapping the one-to-one orthologous gene pairs 1,000 times (Supplemental Table S6).

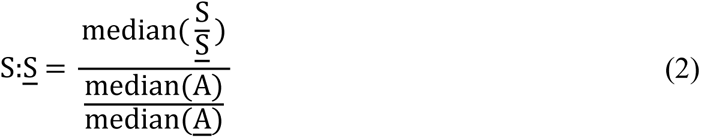

### Expression ratio between ancestral sex-linked genes and ancestral autosomal genes

Similarly, we applied the filter-by-fraction strategy to choose the identical fraction of the most highly expressed ancestral sex-linked genes and the most highly expressed ancestral autosomal genes in our analysis. We calculated the <u>S</u>:<u>A</u> ratio as the median expression levels of the ancestral sex-linked genes divided by the median expression levels of the ancestral autosomal genes (see Formula 3 below). The 95% confidence intervals of the <u>S</u>:<u>A</u> ratios were estimated by bootstrapping the ancestral genes 1,000 times.

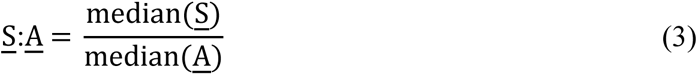

### Expression ratio between current autosomal genes and ancestral autosomal genes

Similarly, we calculated the A:<u>A</u> ratio as the median expression ratio between A and <u>A</u> divided by the ratio of the median expression of autosomal genes in focal species and the median expression of autosomal genes in the out-group (see Formula 4 below). The 95% confidence intervals of these ratios were estimated by bootstrapping the one-to-one orthologous gene pairs 1,000 times.

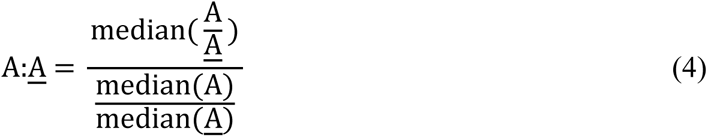

### Estimation of bias in the distribution of housekeeping genes on current and ancestral chromosomes

We used housekeeping genes as a proxy for dosage sensitive genes because (i) identifying housekeeping genes with the expression dataset at hand was easy and accurate; (ii) using the ubiquitously expressed housekeeping genes automatically minimized the potential inaccurate expression measurement due to low expression in some tissues; (iii) the relatively large number of housekeeping genes ensured statistical power in testing the uneven distribution of dosage sensitive genes across multiple species. Following a previously used definition of housekeeping genes (Eisenberg and Levanon 2013), we defined housekeeping genes for a species as those genes with TPM values > 10 in all tissues collected in this study (Supplemental Table S7). We then estimated the housekeeping gene distribution bias between sex chromosomes and autosomes as well as between proto-sex chromosomes and proto-autosomes. For each species, we counted the numbers of housekeeping genes and non-housekeeping genes on sex chromosomes and autosomes, respectively, and assessed the depletion of housekeeping genes on sex chromosomes relative to autosomes by the chi-square test. For comparisons between the proto-sex chromosomes and proto-autosomes, we required that more than 20% of the genes in the ancestral sex chromosome were orthologs to genes in the current sex chromosome. We counted housekeeping genes and non-housekeeping genes on proto-sex chromosomes and proto-autosomes, respectively, and again tested for the depletion of housekeeping genes on proto-sex chromosomes relative to proto-autosomes by the chi-square test.

### Estimation of the S:A ratio at the DNA level

To collect relevant genomic DNA-seq data, we downloaded a compressed SQL file named “SRAmetadb.sqlite” on June 2019, via the R package SRAdb (Zhu, et al. 2013). We used the Perl package “Bio::LITE::Taxonomy::NCBI” to find the taxonomic IDs of each selected species and then further screened the DNA-seq genomic profiles with the following criteria: (i) the genomic profiles were DNA-seq profiles based on the Illumina sequencing platform; (ii) the genomic profiles must have clear sex information; (iii) the genomic profiles were not derived from pathological, stress-treated or genetically modified samples; and (iv) the SRA record had the greatest sequencing depth among records for each sex in each species. If the SRA record could not be downloaded, we discarded the SRA record and re-selected the next best record according to the above rules until all conditions were met. The final genomic dataset consisted of 44 SRA files from 22 species (Supplemental Table S8).

The sex chromosomes to autosomes (S:A) ratio at the DNA level was obtained by dividing the average number of short reads alignable to the sex chromosomes by the average number of short reads alignable to the autosomes. The relative S:A ratio at the DNA level was then calculated by dividing the S:A ratio of the heterogametic sex at the DNA level by the S:A ratio of the homogametic sex (see Formula 5 below). The 95% confidence intervals of these relative ratios were estimated by bootstrapping the short reads over the sex chromosomes and the autosomes 1,000 times.

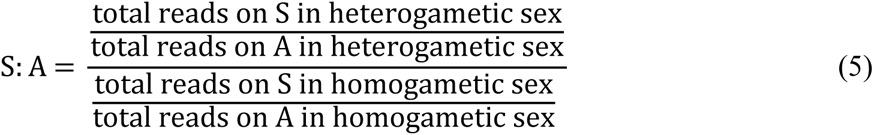

### Phylogenetic independent contrast analysis

Phylogenetic independent contrast was conducted as implemented in “ape” package in R (Paradis, et al. 2004).

## Acknowledgments

We thank Jianfeng Yang for providing computational support; Feng Chen and Xiaoyu Zhang for helpful discussions; and the anonymous reviewers for their insightful and helpful comments on this work. The research was supported by grants from the National Key R&D Program of China (project number 2017YFA0103504 awarded to X. C., and project number 2018ZX10301402 awarded to J.-R. Y.) and the National Natural Science Foundation of China (project numbers 31771406 awarded to X. C., and 31671320, 31871320 and 81830103 awarded to J.-R. Y.).

## Supplemental Figures

**Figure S1.**
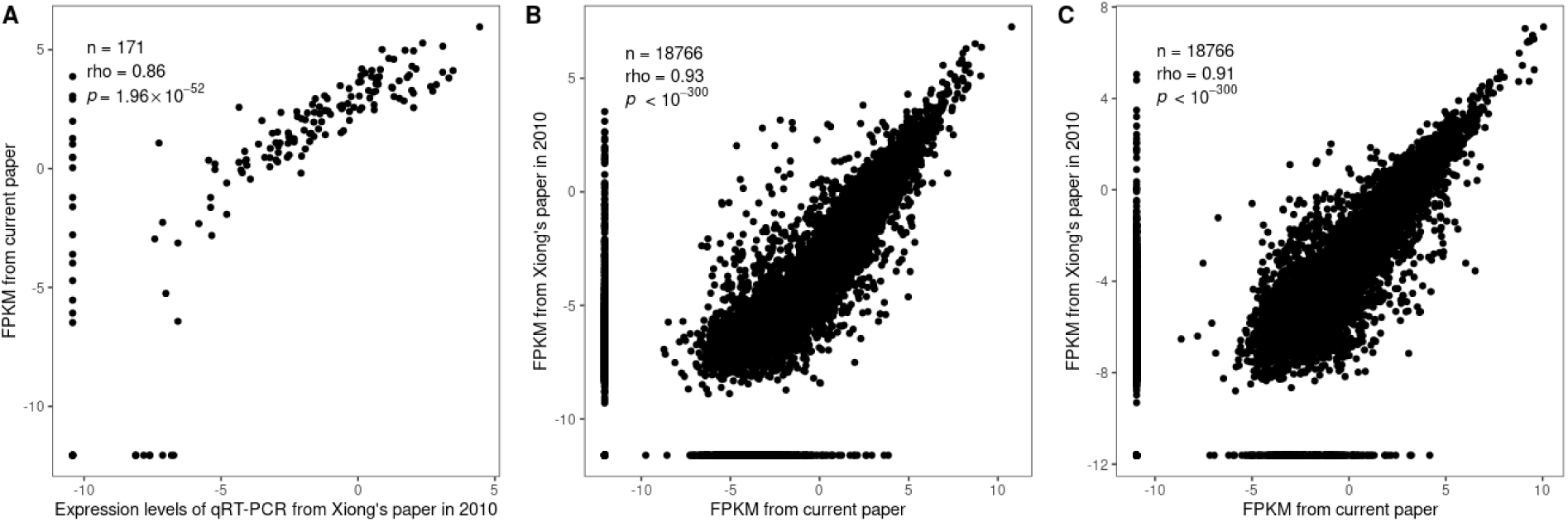
Our pipeline for estimating the transcript abundance in each sample has high accuracy. Our estimated expression levels in male mice were highly correlated with previously reported liver levels determined by qRT-PCR **(a)** and RNA-seq **(b),** with previously reported muscle levels determined by RNA-seq **(c)**. Each dot represents a gene. The FPKM and qRT-PCR expression values were converted with a base-10 logarithmic transformation. The expression abundance of a gene with an FPKM value equal to 0 was set as a tenth of the minimum non-zero expression abundance in each corresponding sample.

**Figure S2.**
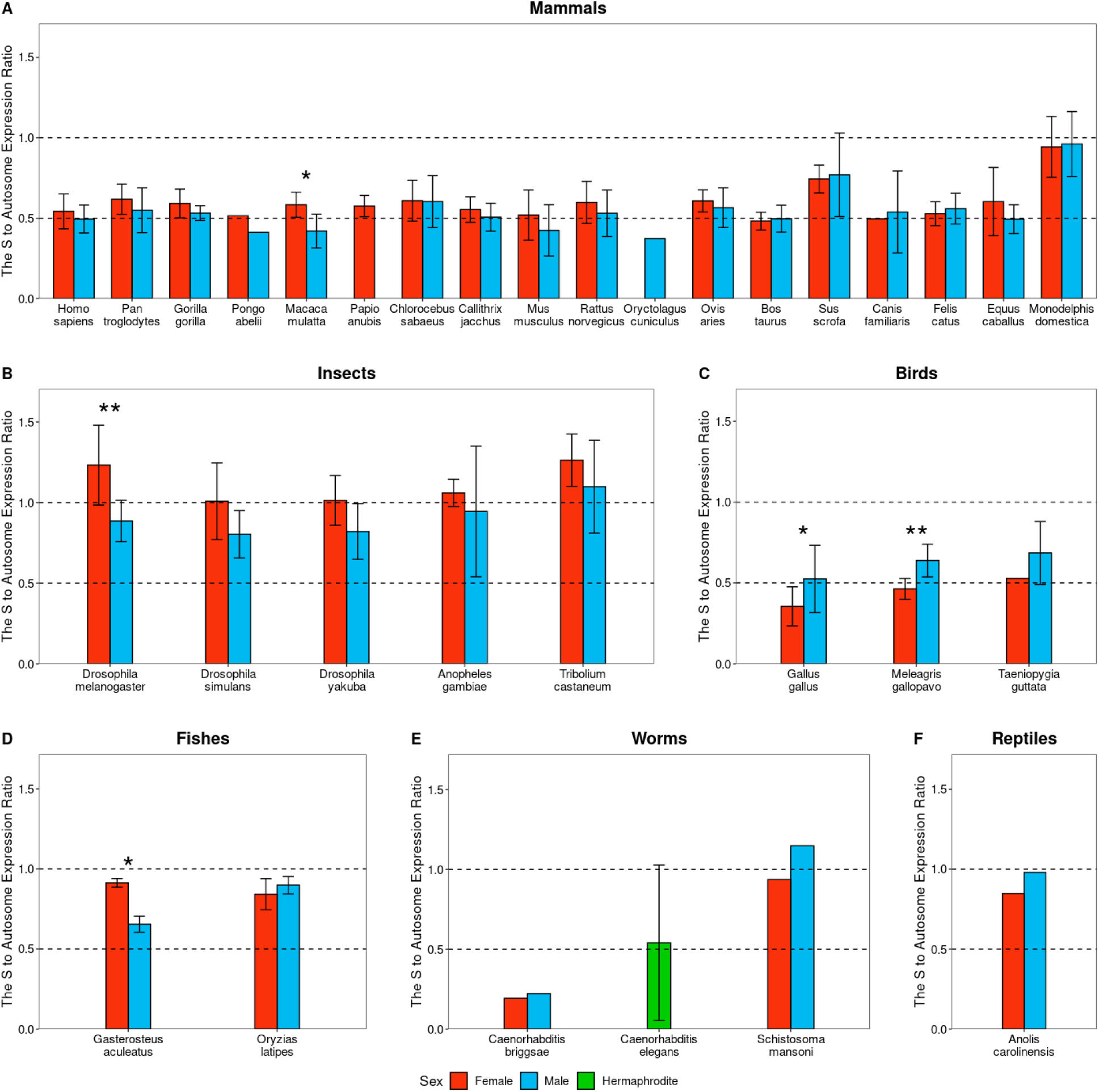
The sex chromosome to autosome expression ratios in multiple species show variations between males and females. The average expression ratios of sex-linked genes and autosomal genes in males and females are shown separately for **(A)** mammals, **(B)** insects, **(C)** birds, **(D)** fishes, **(E)** worms and **(F)** reptiles. The error bars indicate the standard deviations. The red bars represent female tissues, the blue bars represent male tissues, and the green bars represent hermaphrodite tissues. The significance of the difference in the average expression ratios of sex-linked genes and autosomal genes between males and females was tested with Student’s t test (*: *P* < 0.01; **: *P* < 0.001).

**Figure S3.**
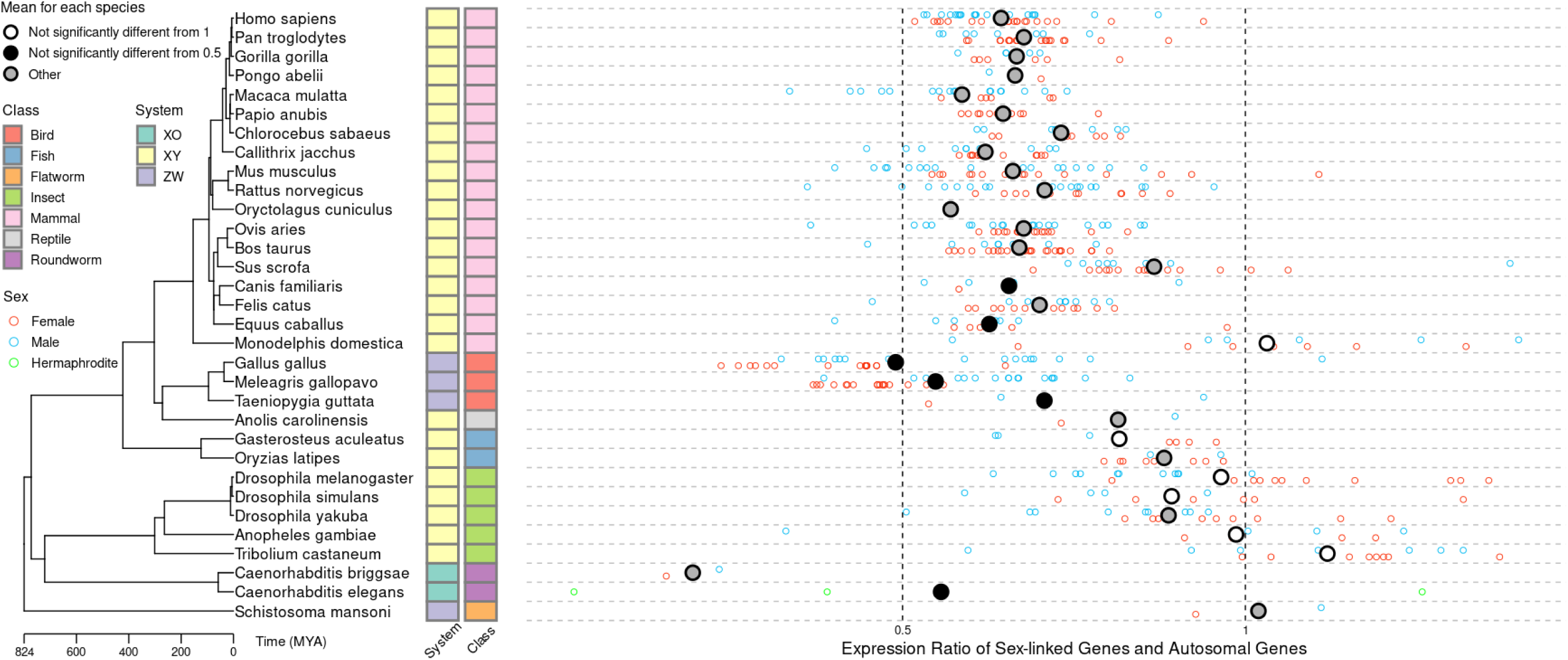
The sex chromosome to autosome expression ratios in multiple species show large variations in the level of sex chromosome dosage compensation. Similar to Fig 2 except that the filter-by-fraction strategy was applied with the threshold of FPKM > 2.

**Figure S4.**
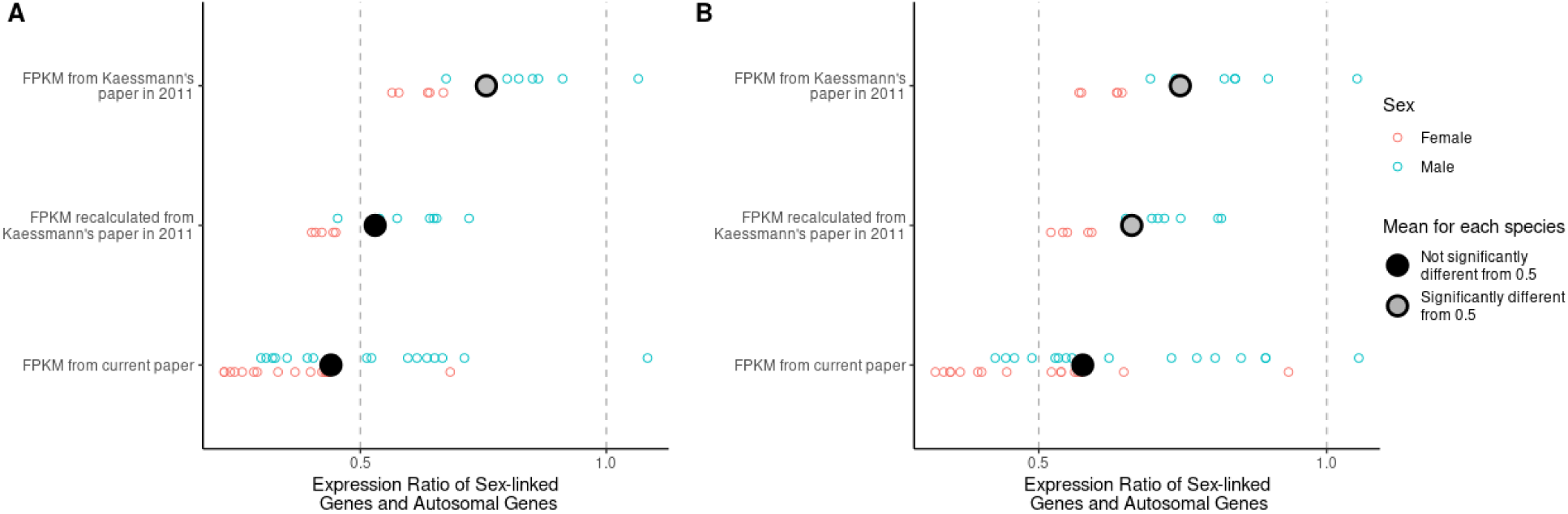
Expression levels derived from our newly compiled dataset are better estimates for the assessment of sex chromosome dosage compensation in birds. The expression ratios of sex-linked and autosomal genes in chickens from the expression levels in the expression levels from Kaessmann’s group, the expression levels from Kaessmann’s group recalculated through our pipeline, and our current dataset are shown. **(A)** The median expression levels of the sex-linked and autosomal genes were calculated by the filter-by-fraction strategy, which involved selection of the fraction of genes whose expression levels exceeded the threshold of FPKM > 0. **(B)** The median expression levels of the sex-linked genes and the autosomal genes were calculated by the filter-by-expression strategy, which involved selection of all genes with expression levels above the threshold of FPKM > 0. The red circles represent the female tissues, and the blue circles represent the male tissues. The average of the median expression ratios is marked with a large circle, and differences were tested with Student’s t test. Values significantly different from 0.5 are marked with gray circles, while values not significantly different from 0.5 are marked with black circles.

**Figure S5.**
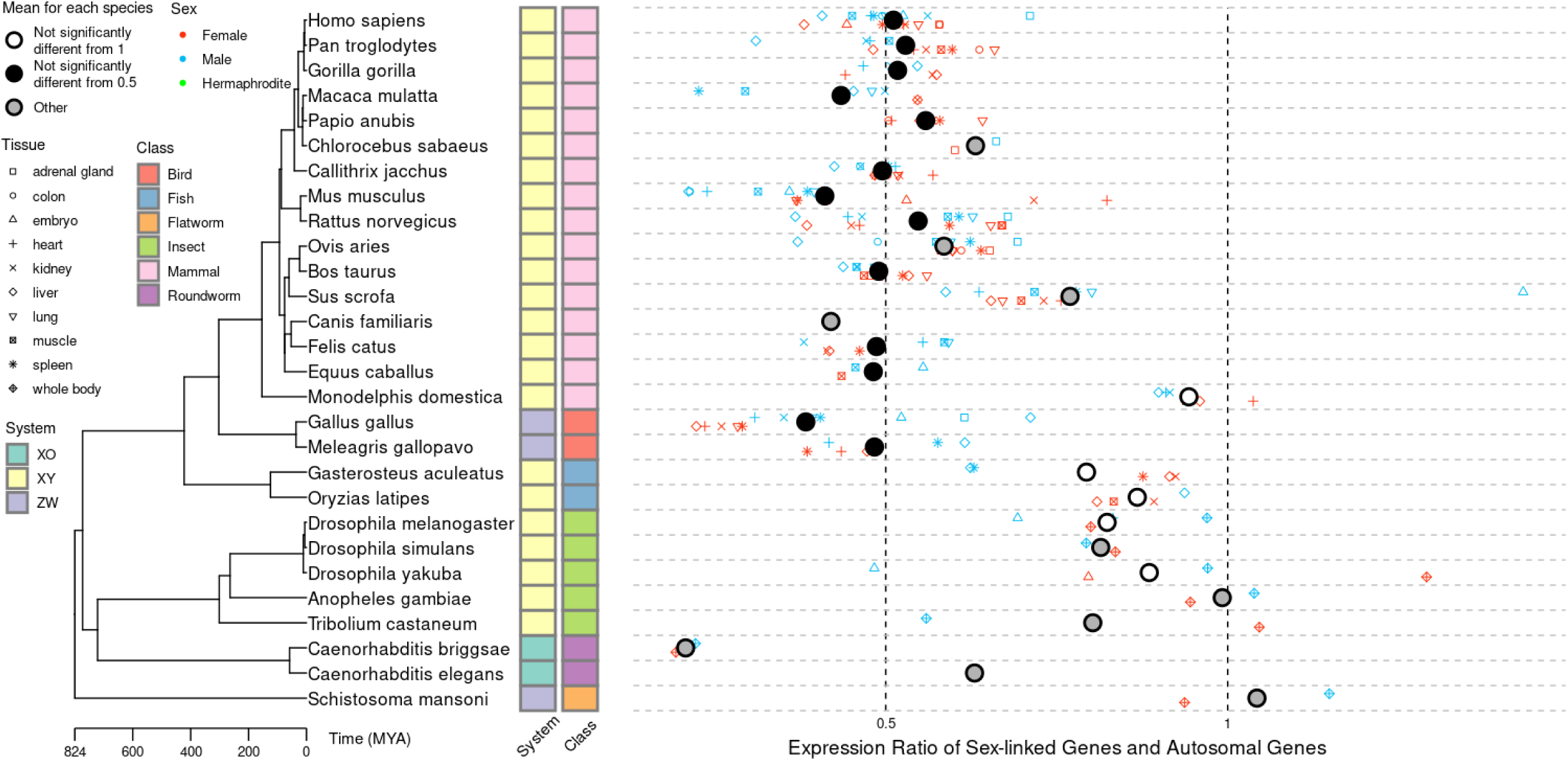
The sex chromosome to autosome expression ratios in multiple species show large variations in the level of sex chromosome dosage compensation. Similar to Fig 2 except that the sex chromosome to autosome expression ratios in ten common (excluding brain and sex-related) tissues labeled by different points of shapes are shown.

**Figure S6.**
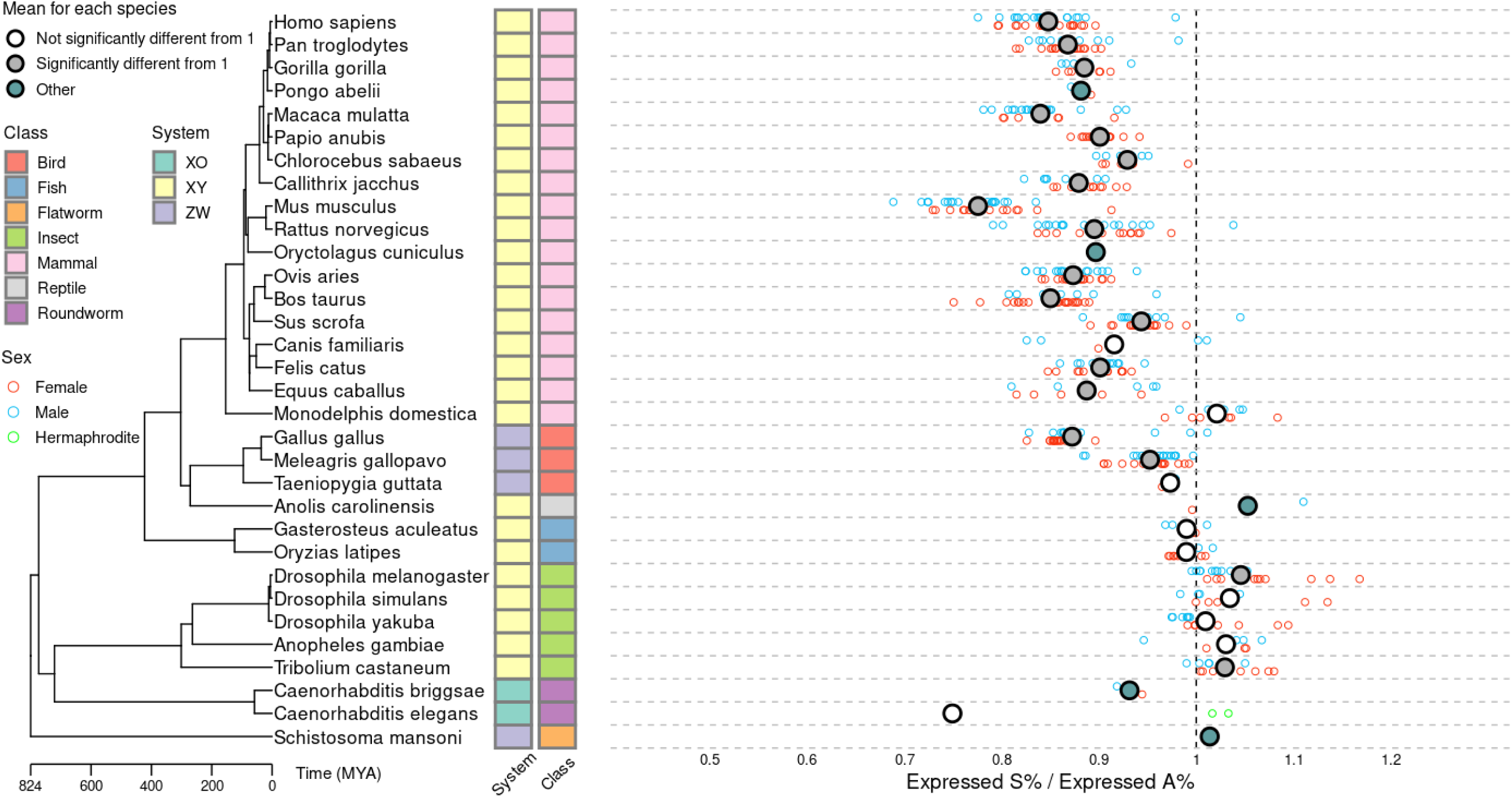
Significant variations in the relative fractions of expressed genes between the sex chromosomes and autosomes in multiple species. Shown on the left is a phylogenetic tree constructed for 32 species whose class and sex systems are labeled with different-colored blocks. The filter-by-expression strategy was applied with a threshold of FPKM > 0 to identify the expressed genes. The fraction of expressed genes on the sex chromosome relative to that on the autosomes for each tissue in each sex of each species is shown on the right. The red circles represent female tissues, the blue circles represent male tissues, and the green circles represent the hermaphrodite tissues. The average fraction of expressed genes on the sex chromosome relative to that on the autosomes in each species is marked with a large circle, and the relative fractions were tested with Student’s t test. Values not significantly different from 1 are marked with white circles, while those significantly different from 1 are marked with gray circles. Average fractions containing less than three corresponding data points (for which Student’s t test could not be performed) are marked with green circles.

**Figure S7.**
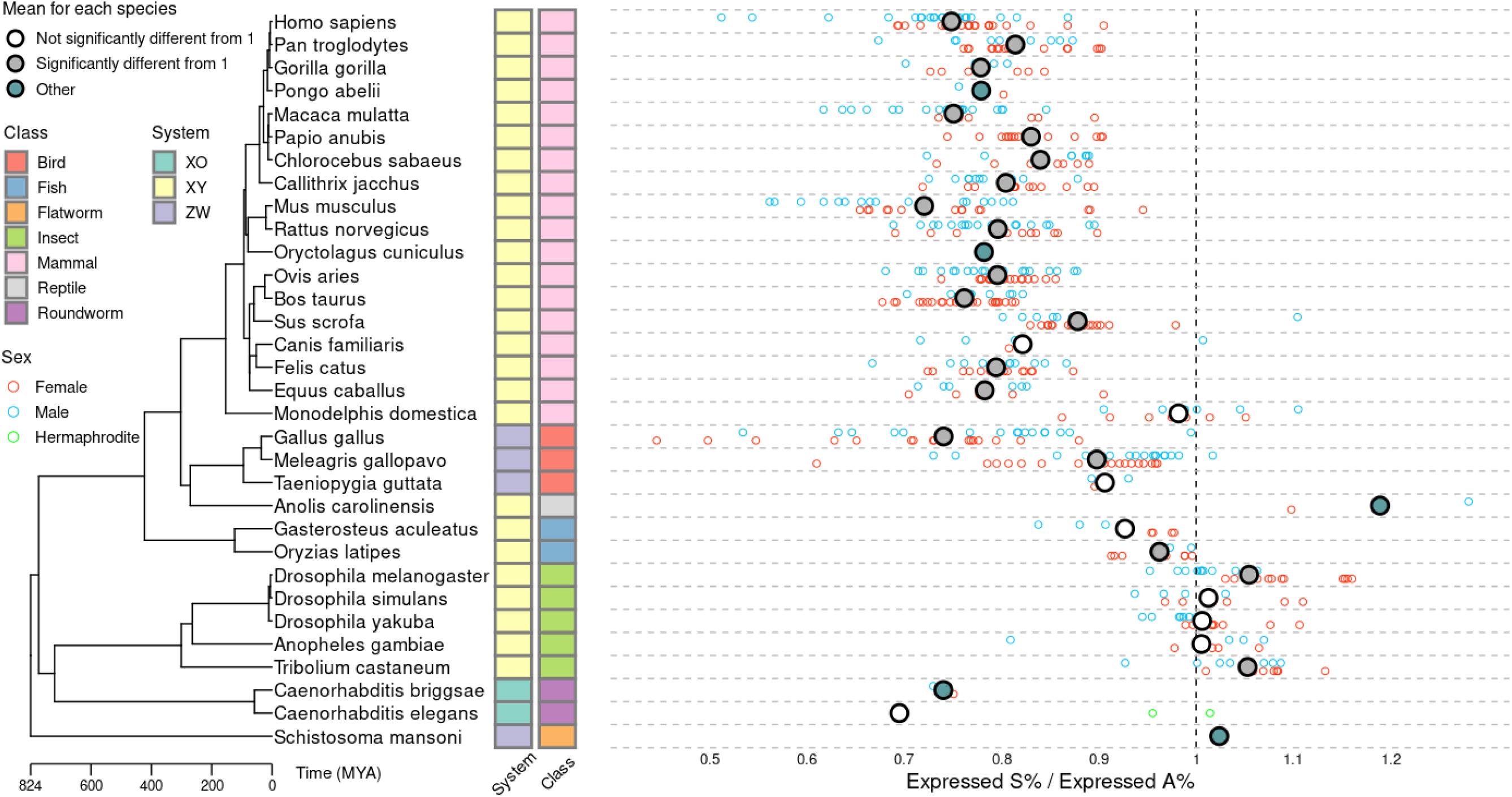
Significant variations in the relative fractions of expressed genes between sex chromosomes and autosomes in multiple species. Similar to Fig S6 except that a filter-by-expression strategy was applied with a threshold of FPKM > 2 to identify expressed genes.

**Figure S8.**
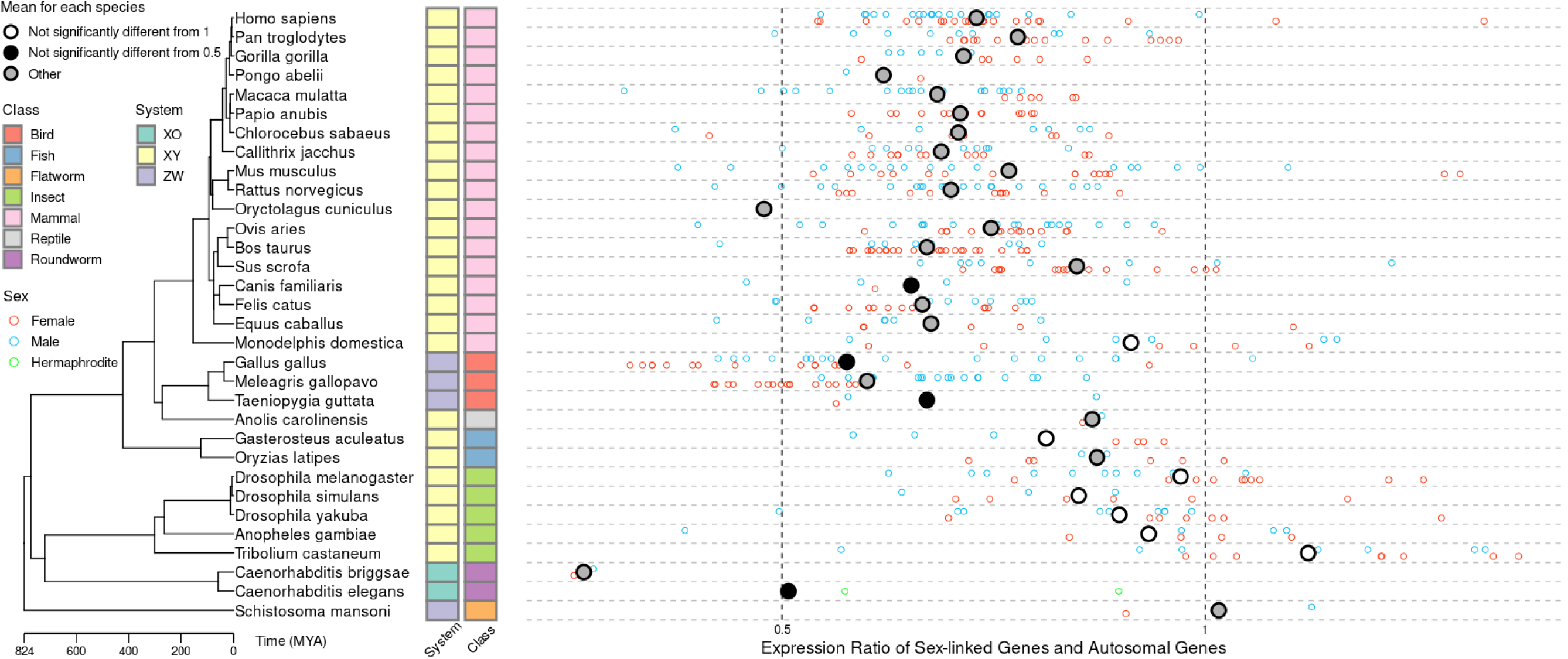
The sex chromosome to autosome expression ratios in multiple species show large variations in the level of sex chromosome dosage compensation. Similar to Fig 2 except that the filter-by-expression strategy was applied with a threshold of FPKM > 0, instead of the filter-by-fraction strategy. The increase in S:A was apparent when compared with Fig 2, especially for the mammals.

**Figure S9.**
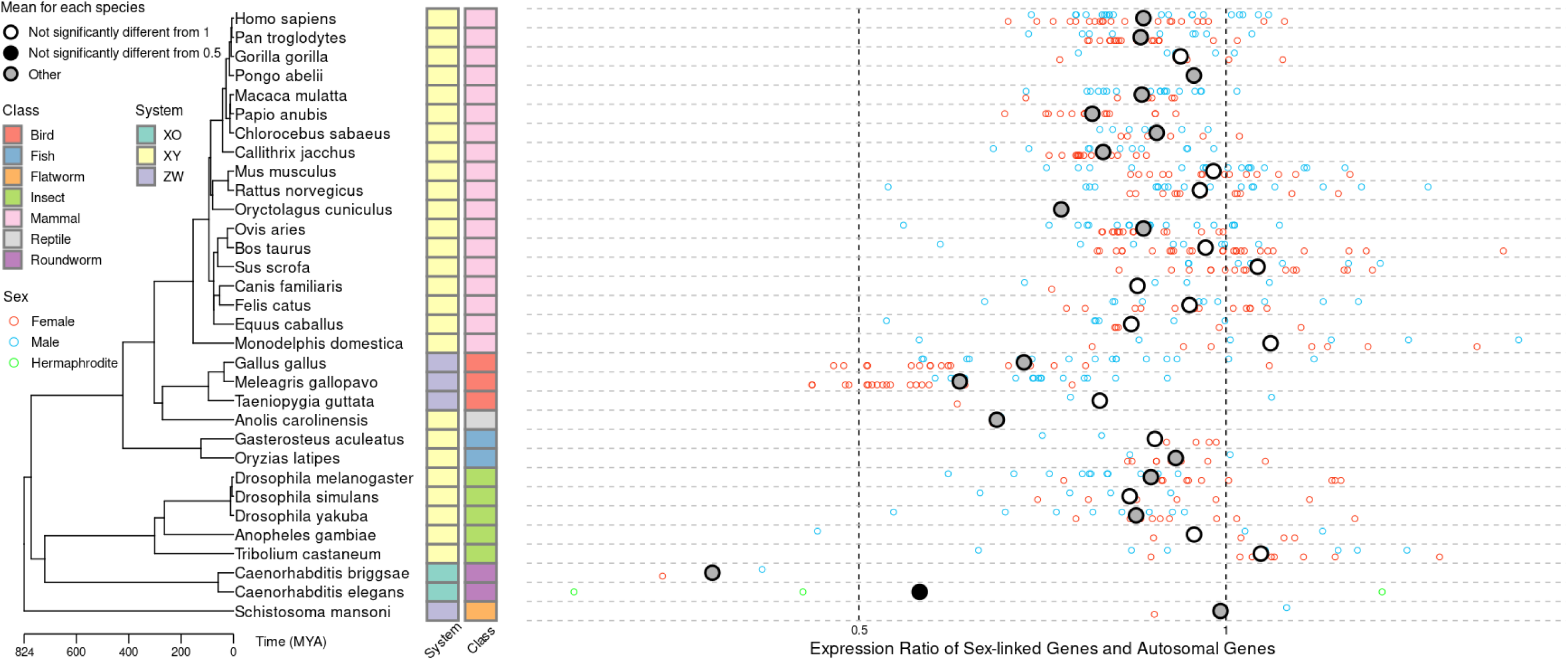
The sex chromosome to autosome expression ratios in multiple species show large variations in the level of sex chromosome dosage compensation. Similar to Fig 2 except that the filter-by-expression strategy was applied with a threshold of FPKM > 2, instead of the filter-by-fraction strategy. The increase in S:A was apparent when compared with Fig 2, and obviously more dramatic than that for Fig S8, especially for the mammals.

**Figure S10.**
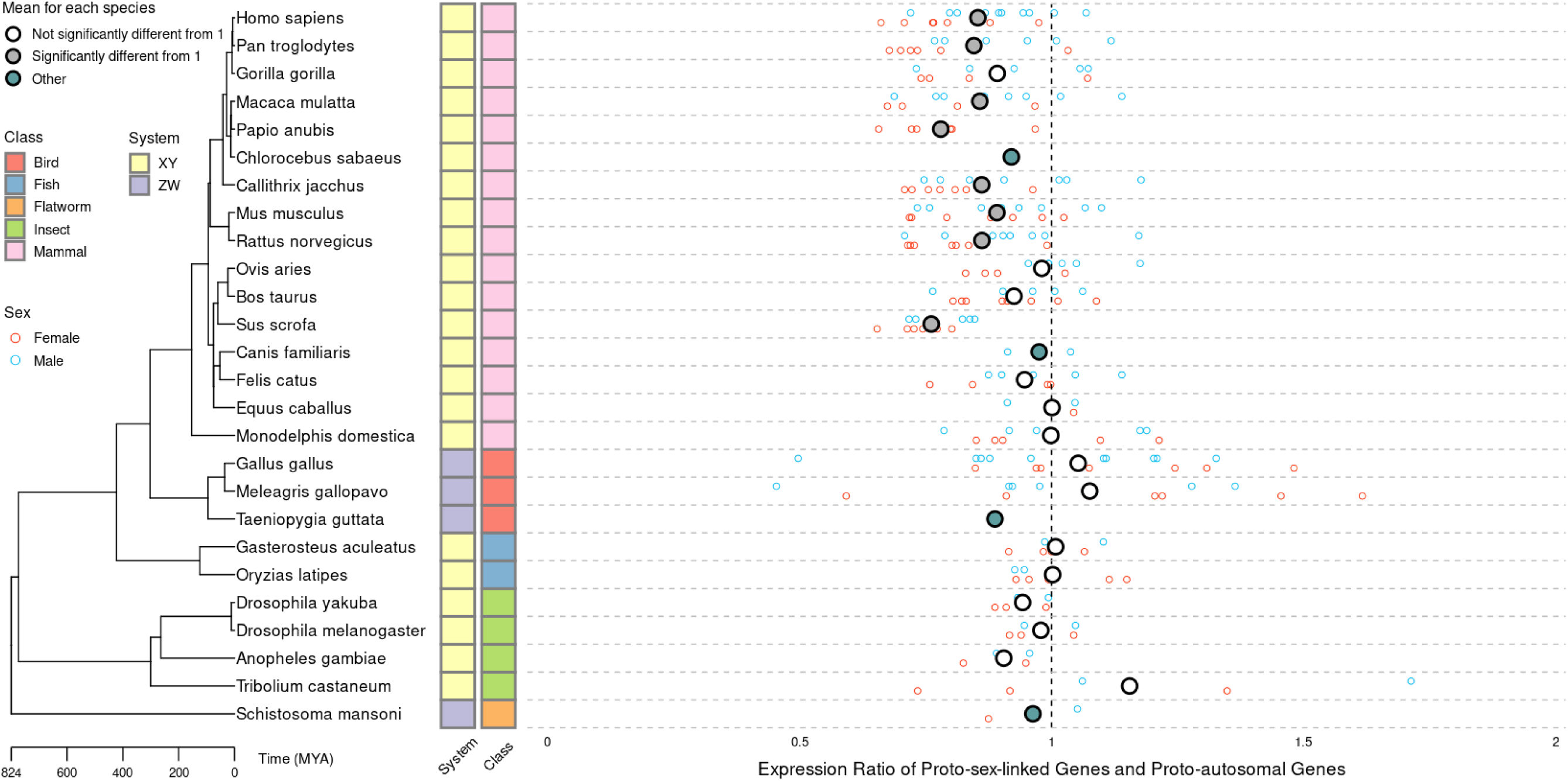
Differences in expression levels between ancestral sex chromosomes and ancestral autosomes in multiple species. Similar to Fig 3 except that the expression ratios of ancestral sex-linked genes and ancestral autosomal genes are shown.

**Figure S11.**
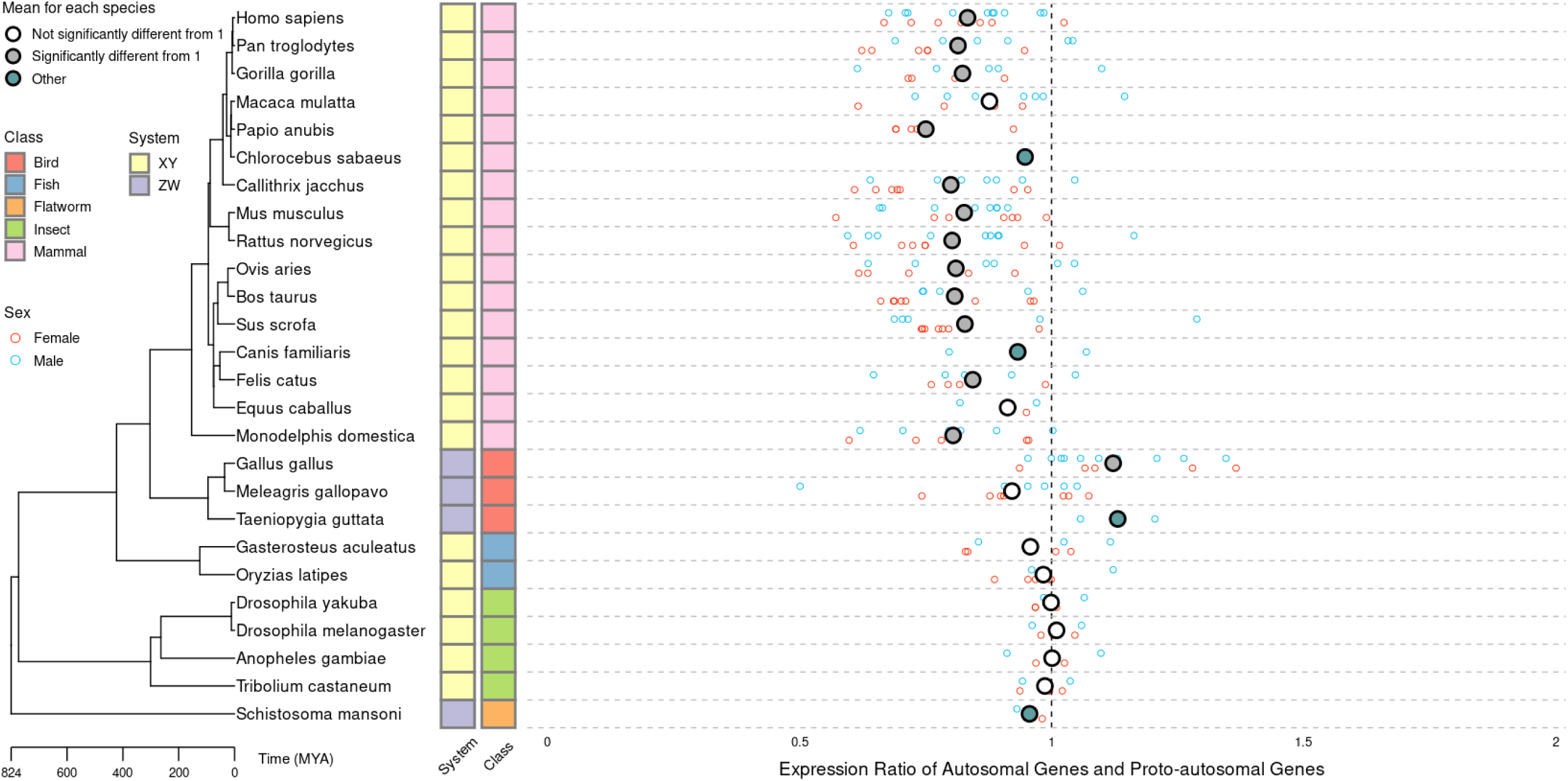
Differences in expression levels between current autosomes and ancestral autosomes in multiple species. Similar to Fig 3 except that the expression ratios of current autosomal genes and ancestral autosomal genes are shown.

**Figure S12.**
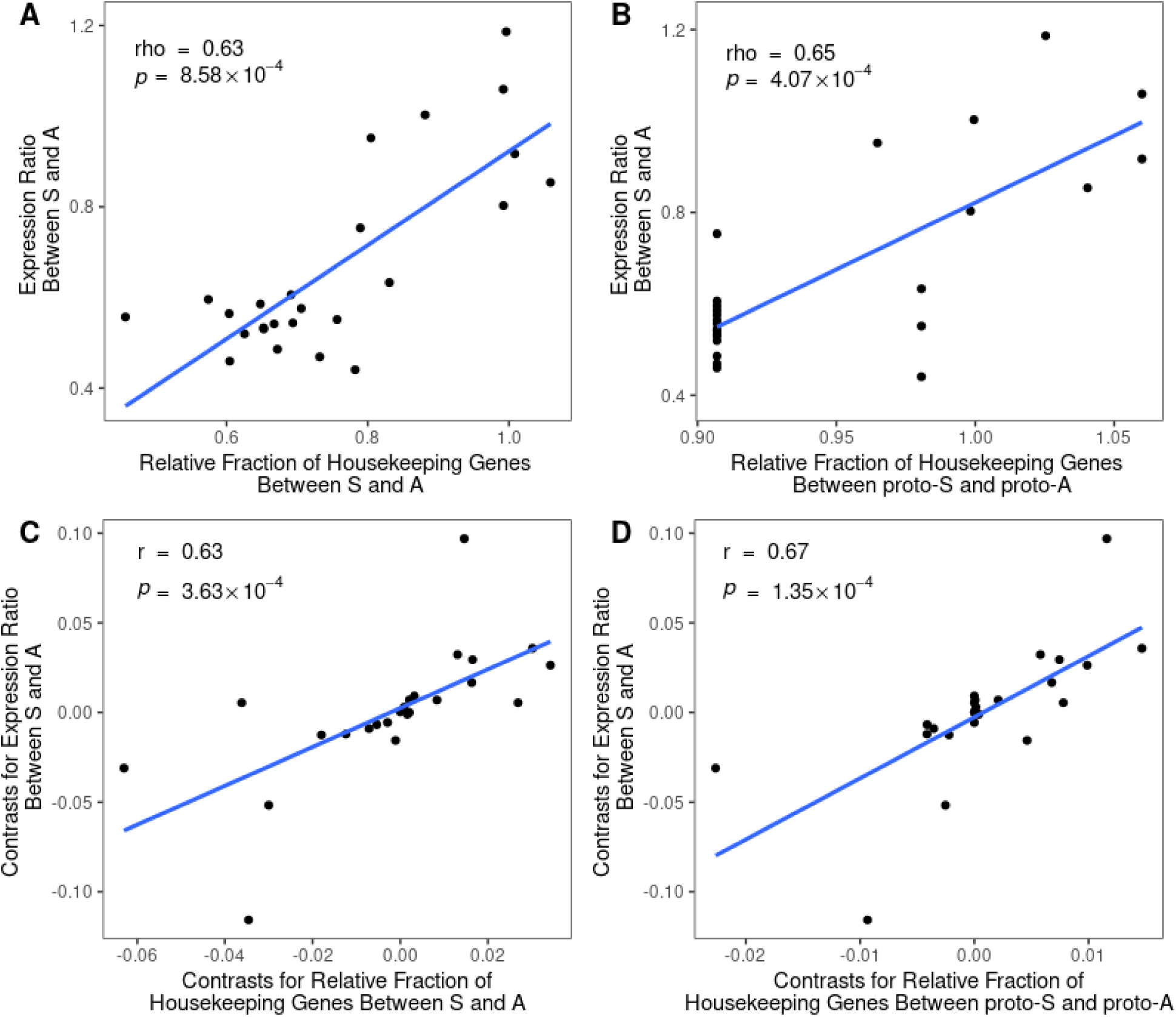
The relative fractions of housekeeping genes were significantly correlated with the sex chromosome to autosome expression ratios. **(A)** and **(B)** are respectively similar to Fig 5A and B, except that the mean S:A ratios in each species are shown. **(C)** and **(D)** are respectively similar to **(A)** and **(B)**, except that the phylogenetic independent contrasts were analyzed for respective variables, thereby correcting the phylogenetic interdependence among them.

